# Bivalent chromatin protects reversibly repressed genes from irreversible silencing

**DOI:** 10.1101/2020.12.02.406751

**Authors:** Dhirendra Kumar, Raja Jothi

## Abstract

Bivalent chromatin is characterized by the simultaneous presence of H3K4me3 and H3K27me3, histone modifications generally associated with transcriptionally active and repressed chromatin, respectively. Prevalent in embryonic stem cells, bivalency is postulated to poise lineage-controlling developmental genes for rapid activation during embryogenesis while maintaining a transcriptionally repressed state in the absence of activation cues, but its function in development and disease remains a mystery. Here we show that bivalency does not poise genes for rapid activation but protects reversibly repressed genes from irreversible silencing. We find that H3K4me3 at bivalent gene promoters—a product of the underlying DNA sequence—persists in nearly all cell types irrespective of gene expression and confers protection from *de novo* DNA methylation. Accordingly, loss of H3K4me3 at bivalent promoters is strongly associated with aberrant hypermethylation and irreversible silencing in adult human cancers. Bivalency may thus represent a distinct regulatory mechanism for maintaining epigenetic plasticity.

**HIGHLIGHTS:**

- Bivalent chromatin does not poise genes for rapid activation
- H3K4me3 at bivalent promoters is not instructive for transcription activation
- H3K4me3 at bivalent promoters protects reversibly repressed genes from *de novo* DNA methylation
- Loss of H3K4me3/bivalency is associated with aberrant DNA hypermethylation in cancer

## INTRODUCTION

The DNA in eukaryotic cells is organized into chromatin consisting of repeating units of nucleosome, an octamer of histone proteins wrapped with approximately 147 base pairs of DNA. Consequently, chromatin plays a central role in regulating accessibility to DNA in many DNA-templated processes including transcription. Histone modifications and DNA methylation are key epigenetic mechanisms that modulate chromatin structure and thus regulate gene expression programs controlling cell fate decisions and cell identity during development (Allis and Jenuwein, 2016; Jaenisch and Bird, 2003; Kouzarides, 2007; Li et al., 2007).

Histones are subject to a vast array of post-translational modifications including acetylation and methylation (Kouzarides, 2007; Li et al., 2007). Whereas histone acetylation is generally associated with gene activation, histone methylation, depending on the residue modified, is associated with either activation or repression. Trimethylation of histone H3 on lysine 4 (H3K4me3) and lysine 27 (H3K27me3) are two of the most extensively studied histone modifications associated with transcriptionally active and repressed chromatin, respectively (Barski et al., 2007). H3K4me3 and H3K27me3, respectively, are catalyzed by the Trithorax group (TrxG) and Polycomb group (PcG) of proteins (Di Croce and Helin, 2013; Margueron and Reinberg, 2011; Piunti and Shilatifard, 2016; Schuettengruber et al., 2017; Simon and Kingston, 2009). Because TrxG and PcG proteins act antagonistically to regulate, respectively, the activated and repressed states of gene expression, H3K4me3 and H3K27me3 were thought to be mutually exclusive. But this assumption was challenged by the discovery of bivalent domains—genomic regions characterized by the simultaneous presence of H3K4me3 and H3K27me3—found predominantly at developmentally regulated gene promoters in embryonic stem cells (ESCs) (Azuara et al., 2006; Bernstein et al., 2006; Blanco et al., 2020; Harikumar and Meshorer, 2015; Shema et al., 2016; Voigt et al., 2012). Although H3K4me3 and H3K27me3 occupy essentially non-overlapping regions within bivalent domains (Barski et al., 2007), with H3K27me3 domains typically flanking a H3K4me3 domain, it was later established that nucleosomes that bear both “active” H3K4me3 and repressive H3K27me3 do exist *in vivo*, albeit on opposite H3 tails in nearly all cases (Shema et al., 2016; Voigt et al., 2012), consistent with direct allosteric inhibition of PRC2 activity by H3K4me3 (Schmitges et al., 2011).

Despite the presence of H3K4me3, bivalently marked promoters are transcriptionally inactive, if not expressed at very low levels (Mikkelsen et al., 2007). This initial observation led to the elegant and inherently appealing concept that bivalency poises/primes lineage-controlling developmental genes for rapid activation during embryogenesis while maintaining a repressed state in the absence of activation cues (Azuara et al., 2006; Bernstein et al., 2006; Voigt et al., 2013); yet, this hypothesis remains to be directly tested, and the function of bivalent domains in development remains a mystery.

Bivalency was initially hypothesized to be an ESC-specific chromatin state. During ESC differentiation, bivalency is thought to resolve into either H3K4me3-only or H3K27me3-only state depending on whether the gene is activated or silenced, respectively. Later observations, however, confirmed the existence of bivalent domains in terminally differentiated cell types (Barski et al., 2007; Mikkelsen et al., 2007; Mohn et al., 2008), raising the question as to its need and functional relevance in cell types with no differentiation potential.

DNA methylation, as an heritable epigenetic mark, adds an additional level of stability by serving as an enduring ‘lock’ to reinforce a previously silenced state by subjecting genes to irreversible transcriptional silencing even in the presence of all of the factors required for their expression (Bestor et al., 2015; Deaton and Bird, 2011; Jones, 2012; Schubeler, 2015). Most gene promoters DNA-hypermethylated in adult human cancers are bivalently marked in ESCs, and it was speculated that bivalency predisposes these genes for aberrant *de novo* DNA methylation and irreversible silencing in cancer (Ohm et al., 2007; Schlesinger et al., 2007; Widschwendter et al., 2007), but evidence supporting this model is largely lacking. Here we set out to decode the function of bivalent chromatin in physiological and pathological settings. Our studies show that bivalent chromatin does not poise genes for rapid activation but protects genes from *de novo* DNA methylation. Genome-wide studies in differentiating ESCs reveal that activation of bivalent genes is no more rapid than that of other transcriptionally silent genes, challenging the premise that H3K4me3 is instructive for transcription. Notably, H3K4me3 at bivalent promoters—a product of the underlying DNA sequence—persists in nearly all cell types irrespective of gene expression and confers protection from de novo DNA methylation. Bivalent genes in ESCs that are frequent targets of aberrant hypermethylation in cancer are particularly strongly associated with loss of H3K4me3/bivalency in cancer. Taken together, our findings suggest that bivalency protects reversibly repressed genes from irreversible silencing and that loss of bivalency makes them more susceptible to aberrant DNA methylation in diseases such as cancer.

## RESULTS

### Bivalent chromatin does not poise genes for rapid activation

To assess whether bivalent chromatin represents a distinct epigenetic state and/or a regulatory mechanism, we investigated genes with bivalently marked promoters in pluripotent human embryonic stem cells (ESCs) (**Table S1; Methods**); henceforth, we use ‘bivalent genes’ to refer to genes whose promoters are bivalently marked in ESCs, unless stated otherwise. To gain insight into the functional significance of bivalent chromatin, we first sought to determine the chromatin and expression status of bivalent genes in lineage-restricted multipotent and terminally differentiated cells. Using publicly available NIH Roadmap Epigenomics project data from a large and diverse collection of human tissues and progenitor cells (http://www.roadmapepigenomics.org/data/), we noted that H3K4me3 enrichment at bivalently marked gene promoters in ESCs persists in nearly all other cell types irrespective of transcriptional activity (**Figure 1A-C; Figure S1A**). In contrast, H3K27me3 enrichment is more dynamic across cell types (**Figure S1B,C**), with its absence not necessarily accompanied by gene activation. Resolution of bivalent state in ESCs to H3K4me3-only state in lineage-restricted cell types coupled with no guarantees of transcription—in the absence of H3K27me3 (**Figure 1A**)—calls into question the premise that the H3K4me3 component of bivalent chromatin poises genes for rapid activation.

**Figure 1.**
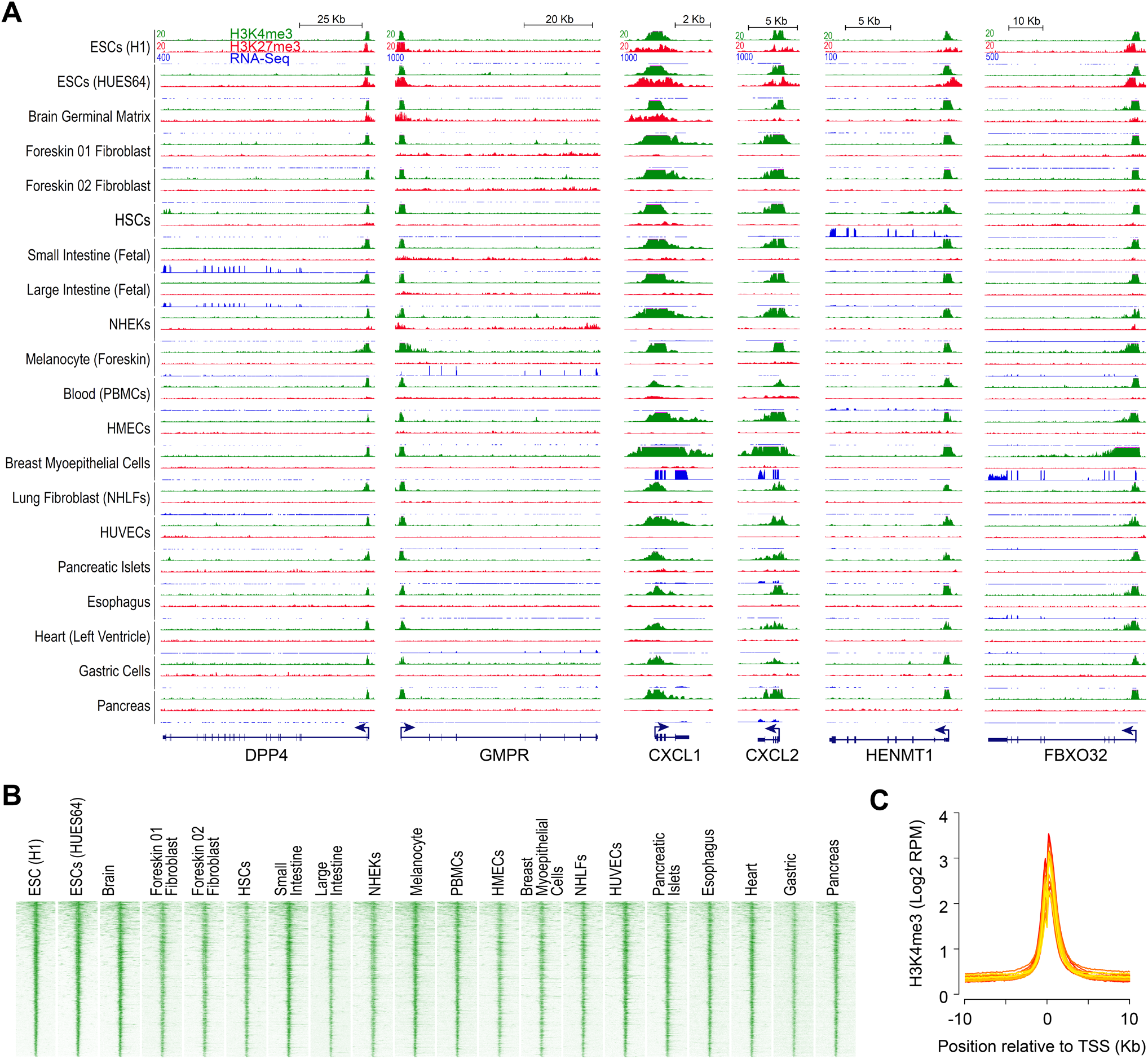
H3K4me3, observed at bivalent promoters in ESCs, persists in nearly all cell types irrespective of gene expression. **(A)** Genome browser shots of select genes, bivalently marked in human embryonic stem cells (ESCs), showing ChIP-Seq read density profiles for H3K4me3 (green) and H3K27me3 (red) in various cell types. Also shown are read density profiles for gene expression (blue; RNA-Seq). HSCs, hematopoietic stem cells; PBMCs, peripheral blood mononuclear cells; HMECs, human mammary epithelial cells; NHEKs, normal human embryonic kidneys, NHLFs, normal human lung fibroblasts; HUVECs, human umbilical vein endothelial cells. **(B)** Heatmap representation of H3K4me3 ChIP-Seq read density, in various cell types, near transcription start sites (TSSs) of genes bivalently marked in human ESCs. Genes were ordered by decreasing order of H3K4me3 signal in ESCs (top to bottom). Read density is represented as RPM (reads per million mapped reads). **(C)** Average H3K4me3 ChIP-Seq read density, in various cell types, near TSSs of genes shown in B. Shades of color represent individual cell types.

To determine whether H3K4me3 at bivalent genes confers them rapid or higher activation potential compared with other transcriptionally silent genes that lack the H3K4me3 mark, we sought to investigate the transcriptional fate of bivalent genes during early embryonic development using a previously validated differentiation system (Buecker et al., 2014; Hayashi et al., 2011; Nakaki et al., 2013; Shirane et al., 2016; Yang et al., 2019) (**Figure 2A**), wherein naïve mouse ESCs—representing the pre-implantation mouse embryo from approximately embryonic day E3.75-E4.5—can be induced to epiblast-like cells (EpiLCs), which most closely resemble the early post-implantation epiblast (E5.5-E6.5). Using the data that we previously generated from chromatin immunoprecipitation sequencing (ChIP-seq) analyses of the chromatin from naïve mouse ESCs using antibodies against histone modifications H3K4me3 and H3K27me3 (Yang et al., 2019), we identified 2,163 genes with bivalently marked promoters (**Figure 2B-D; Table S2; Methods**). For comparison purposes, we also identified genes whose promoters are enriched for H3K4me3 but not H3K27me3 (H3K4me3-only) and vice-versa (H3K27me3-only). Genes with neither H3K4me3 nor H3K27me3 enrichment at their promoters were grouped as ‘unmarked’. Consistent with H3K27me3’s role in maintaining the transcriptionally repressed chromatin state (Riising et al., 2014), nearly all of the bivalent (94%) and H3K27me3-only (99%) genes are transcriptionally inactive, if not expressed at very low levels (<1 FPKM), in ESCs (**Figure 2E**). We note that although bivalent genes are expressed at relatively higher levels (~2-3 fold) compared to transcriptionally silent H3K27me3-only or unmarked genes, their expression is still very low and ~100-fold lower than H3K4me3-only genes, which is consistent with mutual exclusivity between PRC2 activity and active transcription (Riising et al., 2014).

**Figure 2.**
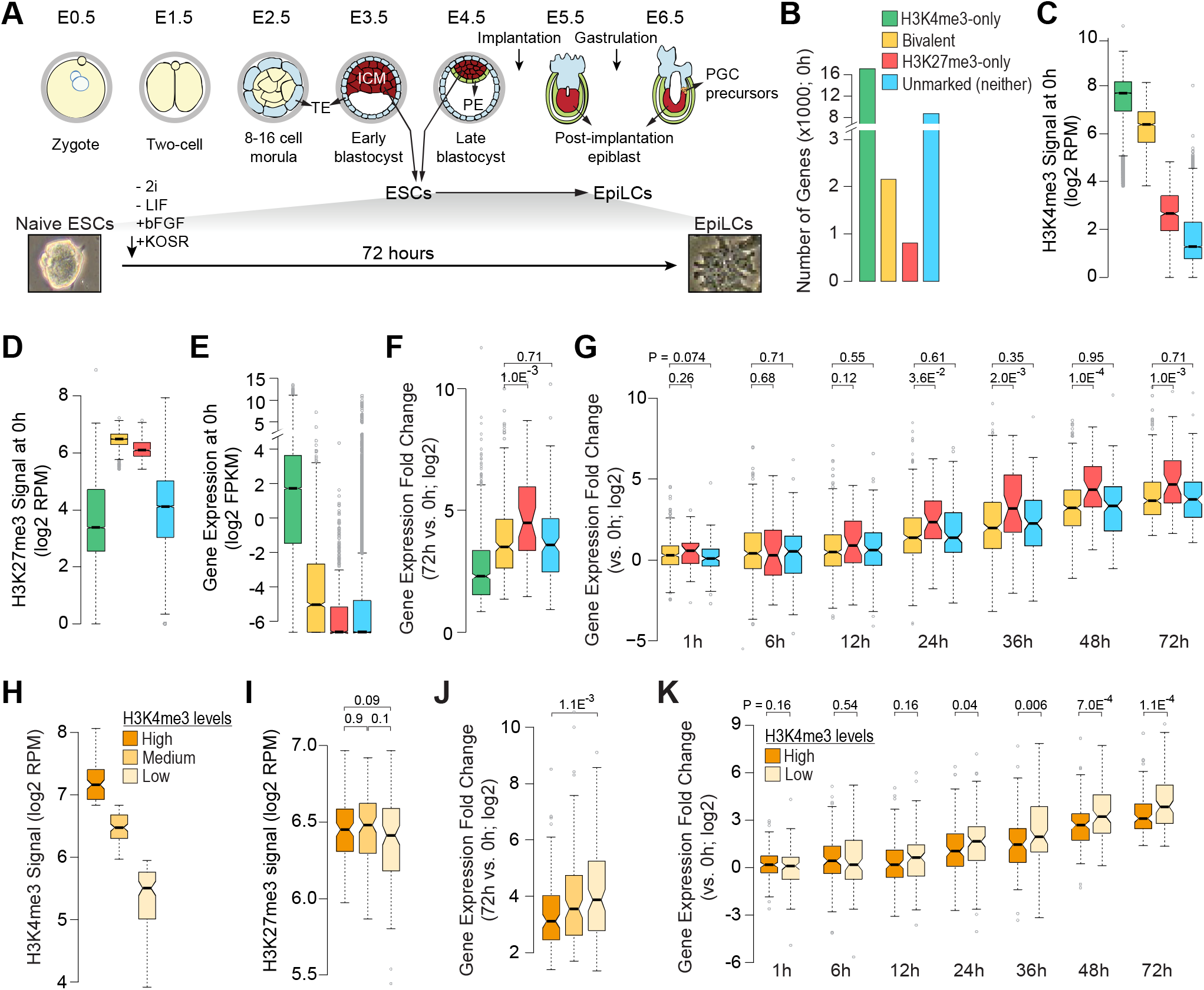
Bivalent chromatin does not poise genes for rapid activation. **(A)** *Top*: Developmental events during early embryogenesis in mouse embryos. *Bottom*: Schematic showing *in vitro* differentiation of naïve ESCs to EpiLCs. ICM, inner cell mass; ESCs, embryonic stem cells; TE, trophectoderm; PE, primitive endoderm; EpiLC, post-implantation epiblast-like cells; PGCs, primordial stem cells. **(B)** Number of genes within each of the four classes, defined based on H3K4me3 (+/-500bp of TSS) and/or H3K27me3 (+/-2Kb of TSS) enrichment at gene promoters in naïve ESCs (0h) (Yang et al., 2019). Bivalent, positive for H3K4me3 and H3K27me3; H3K4me3-only, positive for H3K4me3 and negative for H3K27me3; H3K27me3-only, positive for H3K27me3 and negative for H3K4me3; Unmarked, negative for both H3K4me3 and H3K27me3. **(C-E)** Boxplots showing the distribution of ChIP-Seq read densities for H3K4me3 (**C**) and H3K27me3 (**D**) at gene promoters and gene expression (**E**) in naïve ESCs for each of the four gene classes defined in B. RPM, reads per million mapped reads; FPKM, fragments per kilobase per million mapped reads. **(F)** Boxplot showing the distribution of gene expression fold changes (72h vs 0h) for genes upregulated in EpiLCs (Yang et al., 2019). Genes grouped based on their chromatin states in naïve ESCs (0h). **(G)** Boxplot showing the distribution of gene expression fold changes over time (vs 0h) for genes upregulated in EpiLCs (Yang et al., 2019). Genes grouped based on their chromatin states in naïve ESCs (0h). **(H, I)** Bivalent genes upregulated in EpiLCs (72h vs 0h) were binned into three equal-sized sets based on H3K4me3 enrichment at gene promoters in naïve ESCs (0h). Box plots showing the distribution of ChIP-Seq read densities for H3K4me3 (**H**) and H3K27me3 (**I**) in naïve ESCs for each of the three defined sets. **(J)** Boxplot showing the distribution of gene expression fold changes (72h vs 0h) for bivalent genes upregulated in EpiLCs. Genes grouped based the three sets defined in H. **(K)** Boxplot showing the distribution of gene expression fold changes over time (72h to 0h) for bivalent genes upregulated in EpiLCs (72h vs 0h). Genes grouped based on high/low H3K4me3 signal, as defined in H. All the *P* values were calculated using two-sided Wilcoxon rank-sum test. See also Figure S2.

To evaluate whether bivalent genes are activated any faster or any more than other transcriptionally silent genes (H3K27me3-only or unmarked), we focused on genes upregulated (*q*-value < 0.05) during ESC to EpiLC differentiation (**Table S3**). Our analysis revealed that upregulated bivalent genes are no more activated, measured as either fold change or absolute difference in expression, compared to upregulated H3K27me3-only or unmarked genes (**Figure 2F; Figure S2A**). Next, to address whether bivalent chromatin confers rapid activation potential, we examined gene expression changes at various time points (0, 1, 6, 12, 24, 36, 48, and 72h) during ESC to EpiLC differentiation. We found that activation of upregulated bivalent genes is no more rapid than that of upregulated H3K27me3-only or unmarked genes (**Figure 2G; Figure S2B,C**), challenging the notion that H3K4me3 at bivalent promoters poises them for rapid activation. Nonetheless, if it is true that H3K4me3 at bivalent genes poises them for rapid activation, we reasoned that bivalent genes with higher levels of H3K4me3 must be activated much sooner or much more compared to those with relatively lower levels of H3K4me3. Our analysis of the activation dynamics of upregulated bivalent genes, divided into three equal-sized groups based on their H3K4me3 levels (**Figure 2H,I**), revealed that activation of bivalent genes with higher levels of H3K4me3 is neither greater nor faster compared to those with lower levels of H3K4me3 (**Figure 2K; Figure S2D-F**). Together, these results indicate that bivalent chromatin does not poise genes for rapid activation any more than chromatin marked with just H3K27me3 or chromatin marked with neither H3K4me3 nor H3K27me3.

### Transcriptional competence of ‘poised’ RNA Polymerase II at bivalent genes

Although about two-thirds of the bivalent genes lack transcriptionally engaged RNA Polymerase II (RNAPII) (Williams et al., 2015), the presence of ‘poised’ RNAPII, preferentially phosphorylated at serine 5 but not serine 2 and serine 7, at a subset of bivalent gene promoters has lent some credence to the conceptually appealing notion that bivalency poises genes for rapid activation while keeping them repressed (Brookes et al., 2012; Ferrai et al., 2017; Stock et al., 2007). Phosphorylation of serine 5 on RNAPII, largely mediated by the TFIIH complex, promotes transcription initiation, whereas phosphorylation of serine 2 on RNAPII mediates transition of RNAPII from initiation into productive elongation (Phatnani and Greenleaf, 2006). Interestingly, at a cohort of PRC2-targeted developmental genes, it is Erk2 but not TFIIH that phosphorylates serine 5 on RNAPII, and it was hypothesized that, at these PRC2 target genes, Erk2’s phosphorylation of Serine 5 on RNAPII establishes a poised/stalled form of RNAPII that is competent for transcription (Tee et al., 2014).

To gain further insight into the transcriptional competence of RNAPII observed at bivalent promoters, we focused on bivalent genes that harbor ‘poised’ RNAPII (**Table S4**), defined as RNAPII phosphorylated at serine 5 (S5p^+^) but not serine 2 (S2p^-^) and serine 7 (S7p^-^) (Brookes et al., 2012). We noted that unlike transcriptionally active gene promoters, which are enriched for TFIIH but not Erk2, bivalent gene promoters with ‘poised’ Pol II are enriched for Erk2 but not TFIIH (**Figure 3; Figure S3A**). Consistent with this observation, RNAPII-S5p levels at active promoters correlate with TFIIH (R = 0.62) whereas RNAPII-S5p levels at bivalent promoters correlate with Erk2 (R = 0.5). During early stages of transcription, serine 5 phosphorylation on RNAPII, together with the PAF complex, is known to recruit the histone methyltransferase, Set1/COMPASS, to tri-methylate H3K4 (Shilatifard, 2012), which is reflected in the correlation between RNAPII-S5p and H3K4me3 levels at active genes (R = 0.57). However, RNAPII-S5p levels at bivalent promoters with ‘poised’ Pol II exhibit no such correlation with H3K4me3 levels (R = 0.12) (**Figure S3B**); instead, RNAPII-S5p levels correlate with repression-associated H3K27me3 levels (R = 0.4), which is in marked contrast to the inverse correlation observed between RNAPII-S5p and H3K27me3 at transcriptionally active genes (R = -0.26), but is similar to the correlation observed between RNAPII-S5p and H3K27me3 at H3K27me3-only genes with ‘poised’ RNAPII (R = 0.55). Taken together, these data indicate that the Erk2-mediated serine 5 phosphorylation on RNAPII at bivalent genes is incompatible with transcription.

**Figure 3.**
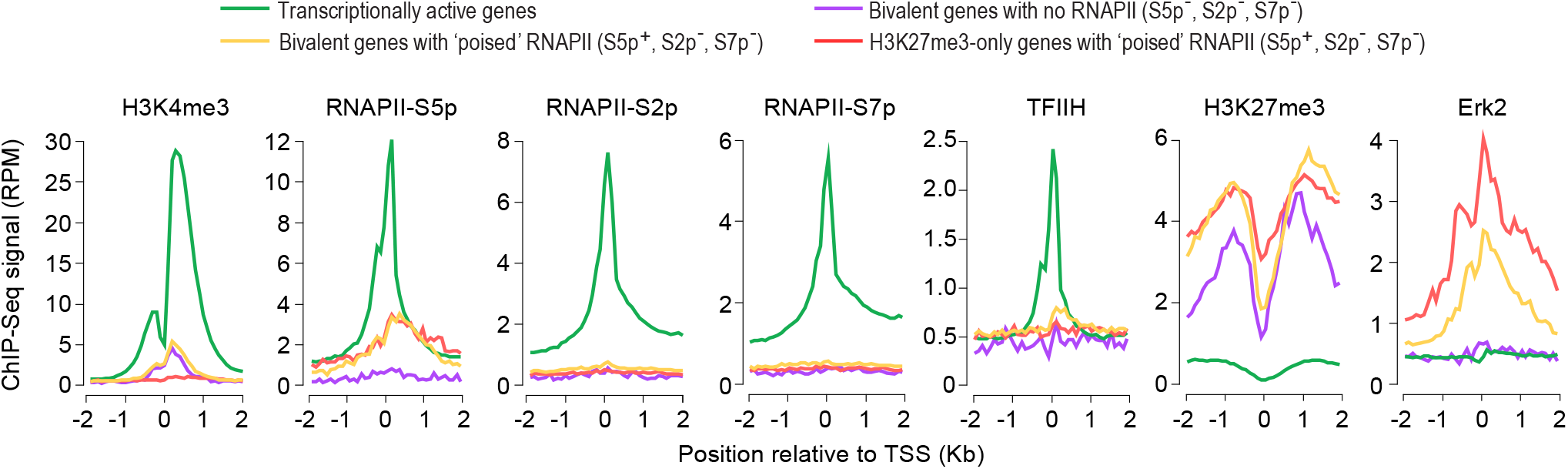
Bivalent promoters with ‘poised’ Pol II are enriched for Erk2 but not TFIIH. ChIP-Seq read density profiles of histone modifications H3K4me3 and H3K27me3 (Marks et al., 2012), phosphorylated forms (S2p, S5p, or S7p) of RNA polymerase II (RNAPII) (Brookes et al., 2012), general transcription factor TFIIH (Ercc3) (Tee et al., 2014) and mitogen-activated protein kinase Erk2 (Tee et al., 2014) near transcription start site (TSS) of indicated gene classes in mouse ESCs grown in serum-containing medium. RPM, reads per million mapped reads. See also Figure S3.

### Bivalent chromatin is a product of PRC2 activity at CpG-rich sequences

Enrichment for developmental genes and regulators among bivalent genes is yet another characteristic that has been used to make the case for the biological relevance of bivalent chromatin in poising lineage-controlling genes for rapid activation during early embryogenesis (Bernstein et al., 2006; Lesch et al., 2016). Although it is true that bivalent genes are enriched for genes associated with developmental processes, this characteristic is not unique to bivalent genes as it also holds true for H3K27me3-only genes (**Figure S4; Table S5**), making it a general feature of genes targeted by PRC2.

With bivalent chromatin conferring no more poising or activation potential than chromatin decorated with just H3K27me3 (**Figure 2G**), we asked why some promoters targeted by PRC2 are also co-enriched for H3K4me3, making them bivalent, while others are not. Because H3K4me3 enrichment at bivalent promoters in ESCs persists in nearly all cell types irrespective of transcriptional activity (**Figure 1**), we reasoned that H3K4me3 at bivalent chromatin is perhaps a product of the underlying DNA sequence features. Indeed, analysis of dinucleotide frequency at H3K27me3-enriched promoters (bivalent and H3K27me3-only) revealed a striking correlation (R = 0.88) between CpG density and H3K4me3 levels (**Figure 4A,B; Table S2**), indicating that CpG density can discriminate between PRC2 targets that are bivalent versus those that are H3K27me3-only. Moreover, this association between CpG density and H3K4me3 levels holds true even for gene promoters that are not PRC2 targets (**Figure 4C**), suggesting that CpG density alone can be an excellent predictor of H3K4me3 levels. A corollary to this conclusion would be that CpG density, together with H3K27me3 levels, can predict bivalent chromatin (**Figure 4B,D**). Indeed, a machine learning approach based on multinomial logistic regression of dinucleotide frequencies and H3K27me3 levels predicted, with >90% accuracy, chromatin status of promoters into one of the four classes (bivalent, H3K27me3-only, H3K4me3-only, or unmarked) (**Figure 4E**). Notably, promoters with bivalent chromatin are predicted with >90% sensitivity (recall) and precision.

**Figure 4.**
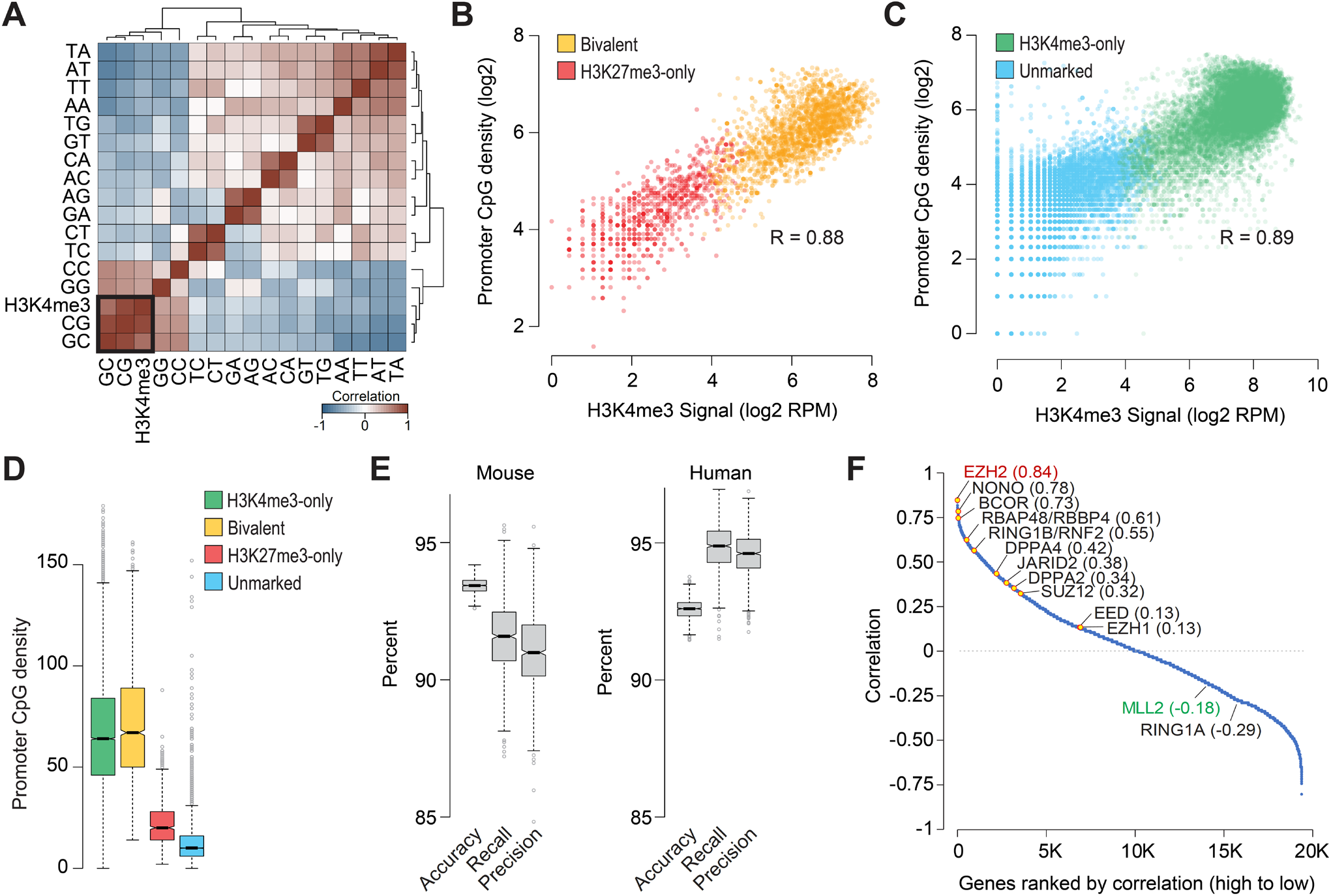
Bivalent chromatin is a product of PRC2 activity at CpG-rich sequences. **(A)** Heatmap showing unsupervised hierarchical clustering of pairwise Pearson correlations between H3K4me3 levels and dinucleotide frequency (5’ to 3’) at the promoters (+/- 500 bp of TSS) of genes enriched for H3K27me3 in naïve mouse ESCs. **(B, C)** Scatter plot showing correlation between CpG dinucleotide frequency and H3K4me3 read density at promoters (+/- 500bp of TSS) of genes with (**B**) or without (**C**) H3K27me3 enrichment in naïve mouse ESCs. **(D)** Boxplots showing the distribution of CpG dinucleotide frequency at promoters (+/- 500 bp of TSS) of genes within each the four classes defined in Figure 2B. **(E)** Statistics summarizing the performance of the multinomial logistic regression-based machine learning method for predicting chromatin state of mouse or human gene promoters (Bivalent, H3K27me3-only, H3K4me3-only, or unmarked) using only H3K27me3 data and dinucleotide frequencies at gene promoters. Boxplots show the distribution of indicated performance measures over 1,000 models. Accuracy, fraction of predictions that are correct; Recall (sensitivity), fraction of bivalent genes correctly predicted as such; Precision (positive predictive value), fraction of predicted bivalent genes that are correct. **(F)** Plot showing Pearson correlation between number of bivalent genes and expression of individual genes, calculated based on data from various cell types. Genes, denoted as individual data points, are sorted (left to right, x-axis) based on their correlation values (y-axis). See also Figure S5.

With DNA sequence features and CpG density in particular seeming to have such an outsized influence on H3K4me3 levels and thus establishment of bivalent domains, we next sought to understand why bivalent domains are more prevalent in pluripotent ESCs compared to terminally differentiated or lineage-restricted multipotent cells (**Figure S5A**). Because bivalently marked promoters across various cell types are CpG-rich (**Figure S5B**) but not *vice versa* (**Figure 4B-D**) and because they almost always harbor H3K4me3 (**Figure 1; Figure S5C**), we reasoned that the prevalence of bivalent domains in a given cell type is presumably a function of the extent of PRC2 activity and/or targeting. Consistent with this notion, analysis of gene expression across various cell types revealed that the number of bivalent genes in a given cell type correlates the best with expression levels of EZH2 (R = 0.84), the catalytic subunit of the PRC2 complex (**Figure 4F; Table S6**). NONO, which is required for ERK activation and Erk2-mediated serine 5 phosphorylation on RNAPII at a subset of bivalent genes (Ma et al., 2016), and BCOR of the non-canonical PRC1 complex, which is required to maintain the repressed transcriptional state of a subset of bivalent genes (Beguelin et al., 2016), are ranked close behind (**Figure 4F**). Interestingly, expression of Mll2 (Kmt2d), the catalytic subunit of the Set1/MLL complex chiefly responsible for H3K4me3 at bivalent domains (Denissov et al., 2014; Hu et al., 2013), correlates negatively (R = -0.18), if at all, with the number of bivalent genes (**Figure 4F; Figure S5D**). Given that CpG-rich sequences, when unmethylated, *per se* are sufficient to establish H3K4me3 domains (Thomson et al., 2010) and that Ezh2 gain/loss-of-function strongly correlates with number of bivalent genes (Shema et al., 2016), these data suggest that bivalent domains are a product of PRC2 activity at genomic regions with high CpG density.

### Bivalent chromatin protects promoters from *de novo* DNA methylation

Gene promoters targeted by PRC2 in ESCs—mostly bivalent genes—are often found to be DNA-hypermethylated in adult human cancers, and it was speculated that bivalent chromatin and/or the presence of Polycomb proteins might predispose these genes for aberrant *de novo* DNA methylation and irreversible silencing in cancer (Ohm et al., 2007; Schlesinger et al., 2007; Widschwendter et al., 2007). Acquisition of promoter DNA methylation at these genes is thought to lock in stem cell phenotypes—at the expense of ability to respond to appropriate lineage commitment and differentiation cues—and initiate abnormal clonal expansion and thereby predispose to cancer (Dawson and Kouzarides, 2012; Easwaran et al., 2014; Jones and Baylin, 2007; Widschwendter et al., 2007).

To determine whether bivalent genes in ESCs are more susceptible to *de novo* DNA methylation during normal development, we investigated promoter DNA methylation levels in naïve mouse ESCs and EpiLCs. ESCs represent the pre-implantation epiblast (~E3.75-E4.5), and EpiLCs most closely resemble early post-implantation epiblast (E5.5-E6.5) (Hayashi et al., 2011; Yang et al., 2019) (**Figure 2A**). This time period spanning pre-to post-implantation epiblast differentiation during early embryonic development is noteworthy because naïve ESCs are associated with global DNA hypomethylation, with a major wave of global *de novo* methylation occurring after implantation (~E5.0) (Greenberg and Bourc’his, 2019; Leitch et al., 2013; Okano et al., 1999; Shirane et al., 2016; Smith et al., 2012), when *de novo* methyltransferases Dnmt3a and Dnmt3b—not expressed in naïve ESCs—get induced by about ~500-1000 fold (**Figure S6A**).

Our analysis of DNA methylation and H3K4me3 levels in naïve ESCs revealed that promoters enriched for H3K4me3 are devoid of DNA methylation (**Figure S6B**). Focusing on the 910 genes whose promoters are hypermethylated in EpiLCs compared to naïve ESCs (**Figure S6C,D**), we noted that bivalent genes in ESCs are significantly under-represented among genes hypermethylated in EpiLCs (**Figure 5A; Figure S6E; Table S7**). In contrast, H3K27me3-only genes in ESCs are significantly over-represented among the hypermethylated genes. Although H3K27me3-only genes in ESCs are 29-fold more likely to be hypermethylated compared with bivalent genes, PRC2 targets in ESCs (bivalent and H3K27me3-only), as a group, are still under-represented among hypermethylated genes (**Figure 5A**), indicating that PRC2 targets are less susceptible to *de novo* DNA methylation at least during early embryonic development. Moreover, because genes hypermethylated in EpiLCs are relatively CpG-poor (**Figure S6F**), over-representation of H3K27me3-only genes among the hypermethylated set is more likely a reflection of their CpG-poor promoters—known to undergo extensive and dynamic methylation and de-methylation during normal development (Meissner et al., 2008)—than their being PRC2 targets. Although the generally inverse correlation between CpG density and hypermethylation (**Figure 4D; Figure 5A**) suggests that protection from *de novo* methylation is perhaps a direct function of the local CpG density, studies in mouse have shown that, for most promoters, high CpG density alone cannot account for their unmethylated state *in vivo* (Lienert et al., 2011), underscoring our limited understanding of the mechanisms that protect most CpG island (CGI) promoters from *de novo* methylation.

**Figure 5.**
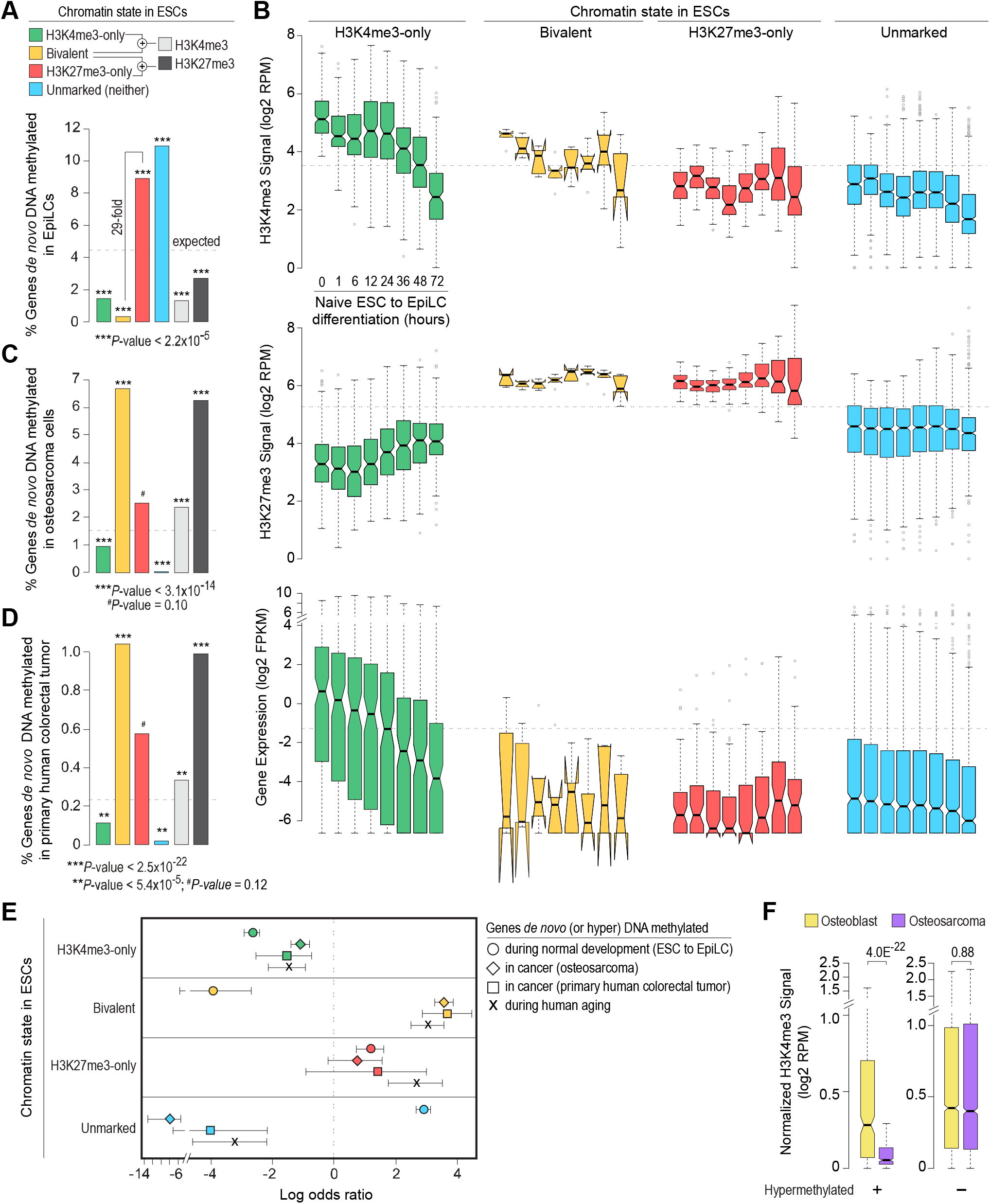
Bivalent chromatin protects promoters from *de novo* DNA methylation. **(A)** Percentage of genes, within each of the four classes of genes defined in naïve mouse ESCs, whose promoters are DNA hypermethylated in EpiLCs (Shirane et al., 2016). **(B)** Boxplots showing changes in promoter H3K4me3/H3K27me3 levels and gene expression (Yang et al., 2019), for each of the four classes of genes shown in A, during ESC to EpiLC differentiation. FPKM, fragments per kilobase per million mapped reads. RPM, reads per million mapped reads. **(C, D)** Percentage of genes, within each of the four classes of genes defined in human ESCs, whose promoters are aberrantly DNA hypermethylated in human osteosarcoma (U2OS) (Easwaran et al., 2012) (**C**) or primary human colorectal tumor (Widschwendter et al., 2007) (**D**). **(E)** Log odds ratio, with 95% confidence intervals, of promoter *de novo* DNA methylation during normal development (mouse ESC to EpiLC differentiation; circle), in cancer (human osteosarcoma and colorectal cancer; diamond and square, respectively), and during aging (Rakyan et al., 2010) (X mark) based on their chromatin state in ESCs. **(F)** Genes bivalently marked in human ESCs were divided into those that are aberrantly DNA hypermethylated in human osteosarcoma (*left*) and those that are not (*right*). Boxplots show the distribution of H3K4me3 levels at these gene promoters in human osteoblasts (yellow) and osteosarcoma (purple) (Easwaran et al., 2012). All the *P* values were calculated using two-sided Wilcoxon rank-sum test. See also Figure S6.

To gain insight into mechanisms that underlie protection of bivalent promoters from *de novo* DNA methylation, we examined chromatin and expression changes that accompany hypermethylation in EpiLCs. High-temporal resolution profiles of H3K4me3, H3K27me3, and gene expression during ESC to EpiLC differentiation (Yang et al., 2019) revealed no major changes in H3K4me3, H3K27me3, or expression levels for hypermethylated H3K27me3-only and unmarked genes (**Figure 5B**), suggesting that *de novo* DNA methylation merely reinforces the previously silenced state at these genes. In contrast, hypermethylated H3K4me3-only genes exhibit a gradual decrease in gene expression and H3K4me3 levels (**Figure 5B**), which would be consistent with the notion that transcriptional activity protects promoters from DNA methylation and that hypermethylation of these genes likely reflects consequence rather than cause of transcription inactivation (Bestor et al., 2015; Jones, 2012; Schubeler, 2015). Although almost all of the bivalent genes in ESCs are protected from *de novo* DNA methylation (**Figure 5A; Figure S6E**), the very few that get hypermethylated in EpiLCs also exhibit a gradual loss of H3K4me3 and thus bivalency, but no change in expression (which was negligible to begin with) or H3K27me3 (**Figure 5B**). Given that *de novo* methylases cannot act on H3K4me3 modified nucleosomes *in vitro* (Ooi et al., 2007) and that complete erasure of H3K4me3 elevates DNA methylation levels (Hu et al., 2009), these findings suggest that H3K4me3 at bivalent promoters protects transcriptionally repressed yet permissive CpG-rich promoters from *de novo* DNA methylation.

### Aberrant DNA methylation in cancer is associated with loss of bivalency

To address whether bivalent chromatin predisposes genes for (or protects genes from) aberrant *de novo* methylation in adult cancer, we analyzed genes hypermethylated in osteosarcoma and colorectal tumor (Easwaran et al., 2012; Widschwendter et al., 2007). About two-thirds of the genes hypermethylated in cancer are bivalent in ESCs (**Figure S6G**). Unlike in EpiLCs, genes bivalent in ESCs are significantly over-represented among genes hypermethylated in cancer (**Figure 5A,C,D**). Importantly, whereas genes that are either H3K4me3-only or H3K27me3-only in ESCs are equally susceptible to hypermethylation during normal development and in cancer, genes bivalent in ESCs are more likely to be hypermethylated in cancer (odds-ratio, OR 11.87, 95% CI: 9.66-14.65 for osteosarcoma, and OR 12.48, 95% CI 7.21-22.34 for colorectal tumor) than during normal development (OR 0.063, 95% CI: 0.02-.15) (**Figure 5E**). To determine whether predisposition of bivalent genes in ESCs to acquire aberrant methylation in cancer might be due to loss of their bivalent status, we examined H3K4me3 and H3K27me3 levels for genes hypermethylated in osteosarcoma. We found that bivalent genes in ESCs that are hypermethylated in osteosarcoma, as opposed to those that are not, exhibit significantly reduced levels of H3K4me3 (**Figure 5F**). No such specificity was observed for H3K27me3 levels; all bivalent genes in ESCs exhibit reduced levels of H3K27me3 irrespective of their methylation status (**Figure S6H**). Altogether, these data suggest that bivalent chromatin protects promoters from *de novo* DNA methylation and irreversible silencing while maintaining a reversibly repressed state, and that loss of H3K4me3 makes these genes more susceptible to aberrant DNA methylation in cancer.

Age is the single biggest risk factor for most diseases including cancer (Niccoli and Partridge, 2012). Because genes frequently hypermethylated and silenced in many adult human cancers exhibit aging-associated hypermethylation and because aging-associated hypermethylation occurs predominantly at promoters bivalently marked in ESCs (Rakyan et al., 2010), we surmised that bivalent genes in ESCs are more susceptible to hypermethylation during the aging process. Indeed, our analysis of aging-associated hypermethylated genes revealed that genes bivalent in ESCs are more likely to be hypermethylated during aging—to the same extent as in cancer—than during ESC to EpiLC differentiation (**Figure 5E; Figure S6I,J**). Moreover, unlike in EpiLCs, promoters of bivalent genes in ESCs that get hypermethylated in cancer and/or during aging are CpG-rich (**Figure S6F,K**). Given that CpG-rich promoters are mostly unmethylated in all cell types at all stages of development, even when transcriptionally inactive (Bestor et al., 2015; Deaton and Bird, 2011; Jones, 2012; Schubeler, 2015), these findings suggest that aging-associated hypermethylation of genes that are bivalently marked in ESCs can serve as a potential biomarker for carcinogenesis in the elderly.

### Establishment and fate of bivalent chromatin

During ESC differentiation and embryonic development, bivalent chromatin is postulated to resolve into either H3K4me3-only or H3K27me3-only state depending on whether the gene is activated or silenced, respectively (Azuara et al., 2006; Bernstein et al., 2006; Voigt et al., 2013). To definitively determine the fates of bivalent genes, we examined chromatin states of gene promoters across various cell types. Our analysis revealed that a vast majority (83%) of genes that are bivalent in ESCs retain H3K4me3 in other cell types, with 52% resolving into H3K4me3-only chromatin state and 31% remaining bivalent (**Figure 6A; Table S8**). Only a small fraction (10%) of bivalent genes in ESCs resolve into H3K27me3-only state in other cell types. Because bivalent promoters are CpG-rich (**Figure 4D; Figure S5B**) and because CpG-rich sequences, when unmethylated, are sufficient to establish H3K4me3(Thomson et al., 2010), these data suggest that bivalent genes have a predilection to resolve into their presumably default H3K4me3-only state in the absence of PRC2 activity. Consistent with this conclusion, we find bivalent promoters that resolve into H3K4me3-only state in most cell types are more CpG-rich compared to those that resolve into H3K27me3-only state in most cell types (**Figure 6B; Table S8**).

**Figure 6.**
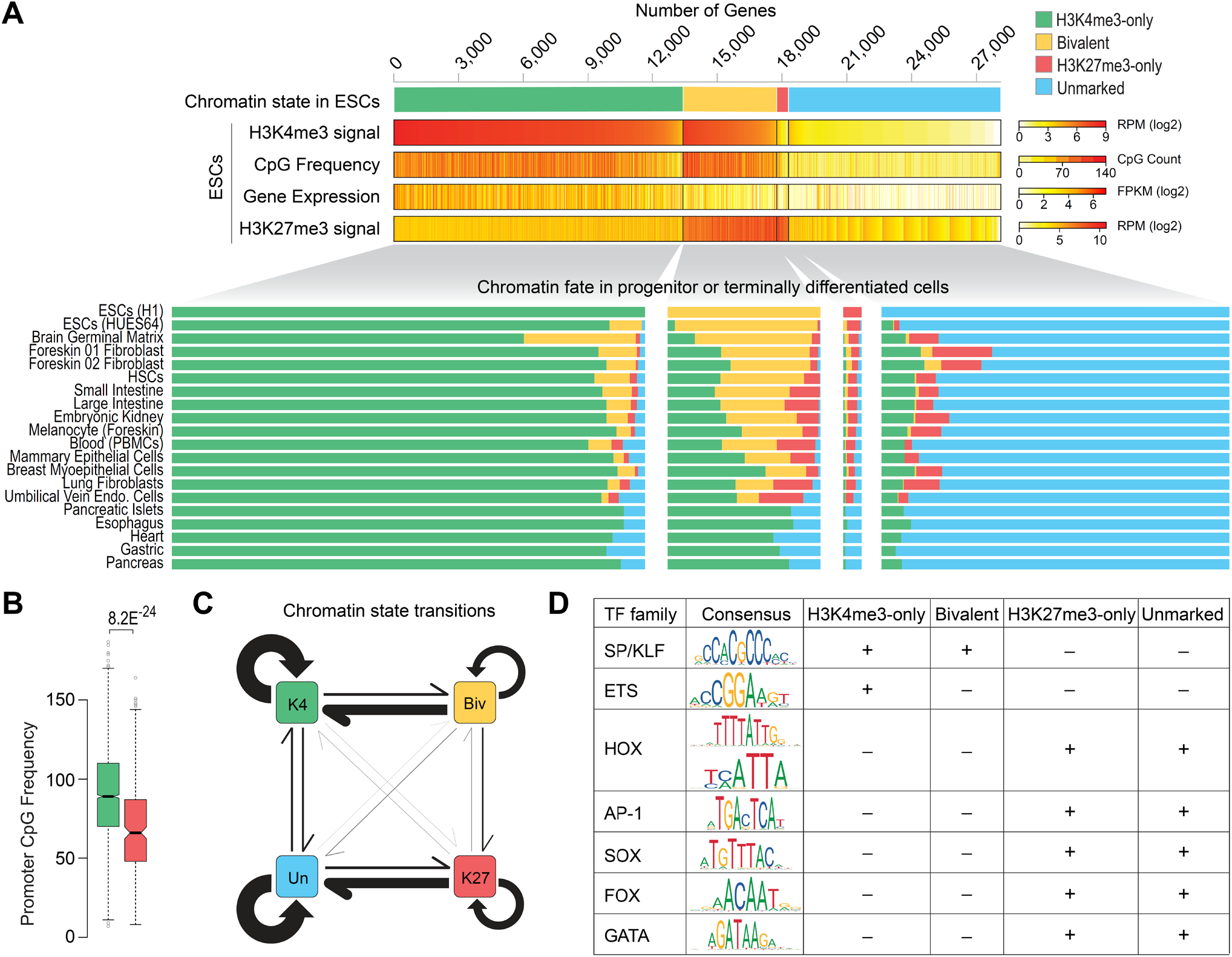
Chromatin fate and sequence characteristics of bivalent promoters. **(A)** *Top*: Genes are grouped into four classes based on their chromatin state, defined based on H3K4me3 (+/- 500 bp of TSS) and/or H3K27me3 (+/- 2 Kb of TSS) enrichment at gene promoters in human ESCs. H3K4me3 and H3K27me3 levels, CpG density (+/- 500 bp of TSS), and gene expression are shown. *Bottom*: Chromatin states of the same four gene classes in various cell types. **(B)** Boxplot showing distribution of CpG density at promoters of bivalent genes (in ESCs) that predominantly resolve into H3K4me3-only (green, left) or H3K27me3-only (red, right) state in other cell types. *P* value calculated using two-sided Wilcoxon rank-sum test. **(C)** Schematic summarizing chromatin state transitions of gene promoters from one state to another. Arrows represent state transitions. The thicker the arrow, the more frequently observed that transition is. K4, H3K4me3-only; Biv, bivalent; K27, H3K27me3-only; Un, unmarked. **(D)** Binding motifs for various transcription factor (TF) families and their enrichment (+) or not (-) within promoters (+/- 500 bp of TSS) of the four genes classes defined in A. See also Figure S7.

Next, to understand the establishment of the bivalent chromatin state, we focused on genes that acquire bivalency in cell types with restricted potency and found that an overwhelming majority (81%) of these genes are H3K4me3-only in ESCs and are CpG-rich (**Figure 6A**). Our analysis of chromatin fates during reconstituted pre-to post-implantation epiblast differentiation in mouse revealed similar results (**Figure S7A**). Notably, nearly all the genes that acquire bivalency in EpiLCs were H3K4me3-only previously. These data further support our conclusion that bivalent chromatin is the culmination of PRC2 activity at regions with high CpG density. Our findings are consistent with studies linking EZH2 to the acquisition of over 1,000 new bivalent genes in germinal center B cells, almost all of which were previously H3K4me3-only in resting B cells (Beguelin et al., 2013).

Lastly, to gain further insight into sequence features—besides CpG density—that underlie bivalent promoters, we explored transcription factor (TF) binding motifs over-represented in H3K4me3-only, bivalent, H3K27me3-only and unmarked promoter classes in ESCs. We found motifs for ubiquitously expressed SP/KLF family of TFs generally over-represented (~2-to 5-fold) within mostly CpG-rich H3K4me3-only and bivalent promoters compared with mostly CpG-poor H3K27me3-only and unmarked promoters (Figure 6D; Figure S7B; Table S9). An exception to this is a small fraction of bivalent promoters—relatively CpG poor—that mostly resolve into H3K27me3-only state in other cell types (Figure 6B); they exhibit no such enrichment for a subset of SP/KLF TF motifs (Figure S7C). Unlike CpG-rich promoters, the largely CpG-poor H3K27me3-only and unmarked promoters are characterized by over-representation of motifs recognized by families of TFs that are tissue-specific (e.g., HOX, AP-1, SOX, FOX, and GATA) (Figure 6D; Figure S7B; Table S9). Interestingly, our analysis also revealed that H3K4me3-only but not bivalent promoters are characterized by over-representation of motifs for ETS family of TFs (Figure S7B), known to activate genes associated with variety of cellular house-keeping processes including cell cycle control, cell proliferation, and cellular differentiation. These data suggest that CpG-rich promoters that are enriched for motifs for SP/KLF but not ETS factors, when transcriptionally inactive, provide a fertile ground for PRC2 activity and establishment of bivalent chromatin, consistent with a causal role for GC-rich sequences—lacking activating TF motifs—in PRC2 recruitment (Mendenhall et al., 2010).

## DISCUSSION

Bivalent genes, by virtue of their exhibiting features of both transcriptionally active and repressed chromatin, are posited as being in a poised state—enabling them to be rapidly activated upon appropriate activation cues during development—while maintaining a transcriptionally repressed state (Azuara et al., 2006; Bernstein et al., 2006; Voigt et al., 2013).

Collectively, our studies reveal that bivalency does not poise genes for rapid activation but protects promoters from *de novo* DNA methylation. Activation of bivalent genes is neither greater nor more rapid than that of other transcriptionally silent genes that lack H3K4me3 at their promoters (**Figure 2G**), challenging the premise that H3K4me3 at bivalent promoters is instructive for rapid activation of transcription. Notably, we find that promoter H3K4me3 levels are a product of the underlying CpG-rich DNA sequence, so much so that CpG density alone can predict H3K4me3 levels/enrichment reasonably accurately (**Figure 4B,C**). This likely explains why H3K4me3 at bivalent promoters in one cell type persists in nearly all other cell types irrespective of gene expression (**Figure 1**) and why unmethylated CGI promoters harbor H3K4me3 even when transcriptionally inactive (Guenther et al., 2007; Mikkelsen et al., 2007). Our findings are consistent with studies showing that high CpG-rich sequences, when unmethylated, *per se* are sufficient to establish H3K4me3 domains (Thomson et al., 2010)—even in the absence of RNAPII and sequence specific TFs (Vastenhouw et al., 2010)— but insufficient to induce transcriptional activity on chromatin (Hartl et al., 2019). Because bivalently marked promoters across various cell types overlap CpG-rich sequences (**Figure S5B**), which inherently are devoid of DNA methylation (Deaton and Bird, 2011) and almost always positive for H3K4me3 (**Figure S5D**), establishment (as well as dissolution) of bivalent domains seems to largely boil down to PRC2 activity (inactivity, respectively) at genomic regions with high CpG density. Supporting this notion, the number of bivalent genes in a given cell type strongly correlate with EZH2 expression (catalytic subunit of PRC2) (**Figure 4F**), with gain- or loss-of-function EZH2 mutation, respectively, associated with increased or decreased bivalency (Shema et al., 2016).

H3K4me3 and H3K27me3 are loosely referred to as “activating” and “repressive” marks respectively, but neither has been firmly established to play a causative role in the regulation of gene expression. To the contrary, it was shown that PRC2/H3K27me3 is not required for the initiation of transcriptional repression of its targets, but is only required for the maintenance of the repressed state (Riising et al., 2014). Despite the general correlation between H3K4me3 and gene expression, it remains unclear as to whether H3K4me3 is instructive for transcription (Howe et al., 2017). Our results showing that the mere presence or the extent of H3K4me3 at bivalent genes does not confer an added advantage when it comes to rapid or higher activation potential **(Figure 2)** suggest that H3K4me3 is not instructive for transcription activation. Consistent with this conclusion, deletion of Mll2—chiefly responsible for H3K4me3 at bivalent chromatin—in mouse ESCs resulted in no substantial disruption in the responsiveness of gene activation after retinoic acid treatment despite the almost complete loss of H3K4me3 and concomitant gain of H3K27me3 at bivalent promoters (Denissov et al., 2014; Hu et al., 2013). Our findings are also consistent with studies in yeast demonstrating that loss of H3K4me3 has no effect on the levels of nascent transcription and, conversely, loss of RNAPII has no effect on H3K4me3 levels (Murray et al., 2019). Together, these observations indicate that H3K4me3 is neither instructive for nor informed by transcription.

Besides its ability to predict transcription or chromatin states, the precise role(s) of H3K4me3 still remains elusive (Piunti and Shilatifard, 2016). Our findings suggest that H3K4me3 is a better predictor of unmethylated CpGs than transcriptional activity and may be a general mechanism to maintain the hypomethylated state of CGIs, even when transcriptionally inactive. This would be consistent with studies showing that H3K4me3 repulses *de novo* methyltransferases *in vitro* (Ooi et al., 2007) and that complete erasure of H3K4me3 elevates DNA methylation levels(Hu et al., 2009; Rose and Klose, 2014). About 70% of mammalian promoters overlap with CGIs (Deaton and Bird, 2011). Although CpG dinucleotides are substrates for DNA methyltransferases, few CGI promoters gain methylation—even when transcriptionally inactive—during normal development (Bestor et al., 2015; Deaton and Bird, 2011; Jones, 2012; Schubeler, 2015). Interestingly, however, most genes that are hypermethylated in cancer have CGI promoters and are bivalently marked in ESCs, which led to the proposition that bivalency predisposes them for aberrant *de novo* DNA methylation and irreversible silencing in cancer (Ohm et al., 2007; Schlesinger et al., 2007; Widschwendter et al., 2007). Our studies reveal that bivalency protects promoters from *de novo* methylation during pre- to post-implantation epiblast differentiation and that aberrant hypermethylation in cancer is explained by the loss of H3K4me3/bivalency (**Figure 5**). In other words, it is not the bivalency but the loss of bivalency that seems to make bivalent genes more susceptible to aberrant DNA methylation in diseases such as cancer.

CGI promoters, the superset containing bivalent promoters, are relatively nucleosome-deficient, intrinsically accessible without the need for ATP-dependent nucleosome displacement, and transcriptionally permissive (Ramirez-Carrozzi et al., 2009). So, what keeps transcriptionally repressed bivalent promoters from getting transcribed? Phosphorylation of serine 5 on RNAPII, which gives rise to its initiated form, is largely mediated by the TFIIH complex (Komarnitsky et al., 2000); accordingly, promoters of transcriptionally active genes are characterized by RNAPII-S5p and TFIIH. Subsequent to the action of TFIIH, the P-TEFb complex phosphorylates serine 2 on RNAPII, giving rise to its transcriptionally-engaged elongating form. Most bivalent promoters lack transcriptionally engaged RNAPII (Williams et al., 2015) but harbor what is referred to as ‘poised’ RNAPII (preferentially phosphorylated at serine 5 but not serine 2), and it has been suggested that ‘poised’ RNAPII primes bivalent genes for rapid activation (Brookes et al., 2012; Ferrai et al., 2017; Stock et al., 2007; Tee et al., 2014). Because promoter-proximal pausing of RNAPII is not a common mechanism employed at bivalent genes (Min et al., 2011; Williams et al., 2015), it is less likely that the ‘poised’ RNAPII at bivalent promoters represents some form of paused/stalled RNAPII competent for rapid transcription re-activation. Our analyses reveal that bivalent promoters are devoid of TFIIH and that any serine 5 phosphorylation on RNAPII at bivalent promoters is attributable to Erk2 (**Figure 3)**, known to bind exclusively to a subset of PRC2 targets (Tee et al., 2014). Because Erk2 and TFIIH are mutually exclusive at their target promoters and because RNAPII-S5p levels at bivalent promoters correlate with H3K27me3 levels and not H3K4me3, as at active promoters, it is conceivable that Erk2-mediated phosphorylation of serine 5 on RNAPII (or Erk’s mere presence on chromatin) is refractory to transcription. In this scenario, Erk2 and/or the substrate it modifies on RNAPII (one or more CTD heptad repeats) may antagonize TFIIH, and activation of transcription likely occurs only upon loss of Erk2 binding and/or Erk2-mediated phosphorylation of serine 5 on RNAPII, which may be followed by binding of appropriate transcription factors at promoters and/or enhancers. Further studies are required to ascertain any potential antagonism between Erk2 and TFIIH.

Mll2 is dispensable for maintaining ESC self-renewal, but Mll2 deficiency is embryonic lethal. Mll2 knock-out (KO) mice exhibit growth defects as early as ~E6.5 and die at ~E10.5 (Glaser et al., 2006), suggesting that Mll2 is not required until after implantation, right when Dnmt3a and Dnmt3b get induced to carry out global *de novo* methylation in early post-implantation embryo. Furthermore, *in vitro* differentiation of Mll2 KO ESCs results in impaired embryoid body formation, with many bivalent genes with key functions in embryonic development and differentiation failing to activate or exhibiting delayed activation kinetics (Lubitz et al., 2007; Mas et al., 2018), indicating an essential role for Mll2 during ESC differentiation. Our finding that H3K4me3 at transcriptionally repressed bivalent promoters, catalyzed primarily by Mll2 (Denissov et al., 2014; Hu et al., 2013), confers protection against *de novo* DNA methylation during pre- to post-implantation epiblast differentiation (**Figure 5A,B,E**) suggests that the requirement for Mll2 after implantation—when it is no longer the major H3K4 trimethyltransferase (Glaser et al., 2006)—at least in part might have to do with its role in implementing H3K4me3 at bivalent genes in order to maintain epigenetic plasticity by protecting against *de novo* DNA methylation and thus irreversible silencing. Because ESCs do not express Dnmt3a or Dnmt3b and are DNA hypomethylated, this could perhaps explain why Mll2 is dispensable in mouse ESCs. Moreover, a recent study showed that Mll2—which also is responsible for H3K4me3 at a vast majority of transcriptionally active genes (Denissov et al., 2014)—protects about 2% of Mll2-dependent active genes from Dnmt1-mediated maintenance methylation (Douillet et al., 2020), highlighting Mll2’s multifaceted role in regulating gene expression.

In summary, our findings suggest a unifying model (Figure 7) wherein bivalency maintains epigenetic plasticity by protecting gene promoters from irreversible silencing while maintaining a reversibly repressed state, and that loss of loss of H3K4me3 makes them more susceptible to aberrant DNA methylation in diseases such as cancer.

**Figure 7.**
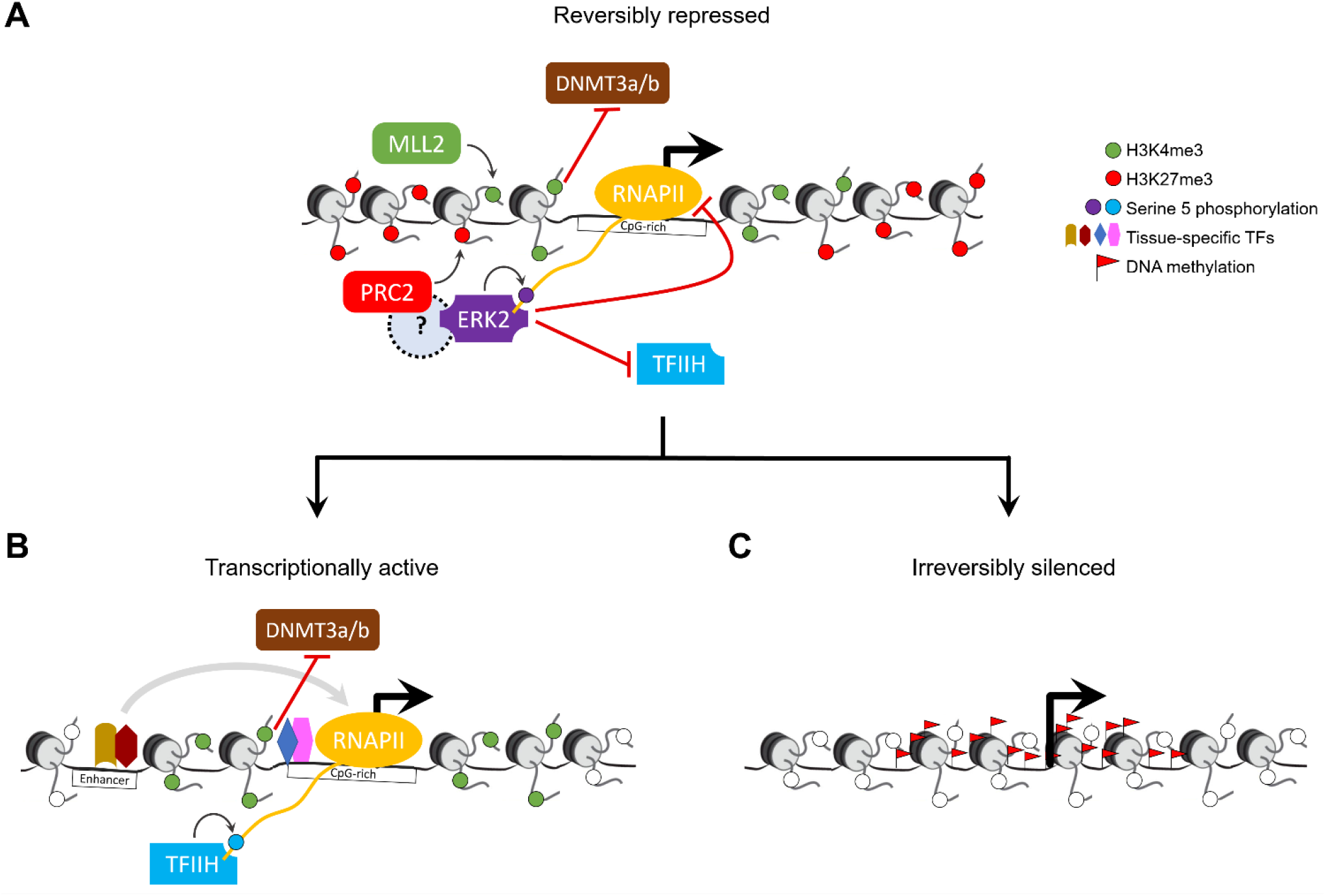
Model for bivalent chromatin maintaining epigenetic plasticity by protecting gene promoters from irreversible silencing while maintaining a reversibly repressed state. **(A)** H3K4me3, catalyzed primarily by the MLL2/COMPASS complex, protects CpG-rich bivalent promoters from DNA methylation by repelling *de novo* methyltransferases Dnmt3a and Dnmt3b. Bivalent promoters, by virtue of their overlapping CGIs, are more averse to assembling into nucleosomes compared to other genomic DNA (Ramirez-Carrozzi et al., 2009); consequently, bivalent promoters are relatively nucleosome-deficient, intrinsically accessible, and transcriptionally permissive (Deaton and Bird, 2011; Mas et al., 2018). The assembly of transcription machinery at the promoter triggers PRC2 to recruit— either directly or indirectly through yet-to-be-determined mechanism—reinforcement in the form of ERK2, which, in lieu of the usual TFIIH, phosphorylates serine 5 on a particular (or a set of) RNAPII CTD heptad repeat(s) (Tee et al., 2014). The presence of ERK2 and/or the ensuing serine 5 phosphorylation is refractory to transcription as it antagonizes the recruitment of the TFIIH complex, which, besides its role in phosphorylating serine 5 on RNAPII, is necessary to unwind promoter DNA to form transcription bubble for RNA synthesis. **(B)** Activation of transcription at bivalent genes likely occurs only upon loss of PRC2 activity and thus ERK2 activity, followed by binding of appropriate transcription factors at promoters and/or enhancers. **(C)** Loss of H3K4me3 at CpG-rich bivalent promoters makes them more susceptible to aberrant DNA methylation during aging and in diseases such as cancer.

## METHODS

### Data sources

#### Mouse

ChIP-Seq datasets for mouse ESC to EpiLC differentiation (H3K4me3, H3K27me3, and corresponding genomic input) (Yang et al., 2019) and mouse ESCs grown in serum-containing medium (H3K4me3 and H3K27me3 (Marks et al., 2012); RNAPII-S5p, RNAPII-S2p, and RNAPII-S7p (Brookes et al., 2012); Erk2 and TFIIH/Ercc3 (Tee et al., 2014)) were obtained from NCBI GEO portal and processed similarly for data consistency. Briefly, single-end reads from the ChIP-seq and genomic input were mapped to the mouse genome (mm9 assembly) using Bowtie (Langmead et al., 2009), allowing for up to three mismatches, retaining only reads that align to unique genomic locations. Processed promoter DNA methylation data from naïve mouse ESCs and EpiLCs were obtained from a previously published study (Shirane et al., 2016). RNA-Seq datasets for mouse ESC to EpiLC differentiation (Yang et al., 2019) and mouse ESCs grown in serum containing medium (Marks et al., 2012) were obtained from NCBI GEO portal and processed the same way for data consistency. List of genes that are (a) transcriptionally active, (b) bivalent and harbor ‘poised’ RNAPII, (c) bivalent but harbor no RNAPII, or (d) H3K27me3-only but harbor poised RNAPII were derived from a previous study (Brookes et al., 2012).

#### Human

Uniformly processed and consolidated Human Epigenome Roadmap data, mapped to the human genome (hg19), for various cell/tissue types were downloaded from the Washington University portal (https://egg2.wustl.edu/roadmap/web_portal/). Only those normal cell/tissue-types for which RNA-Seq, H3K4me3 ChIP-Seq, H3K27me3 ChIP-Seq, and genomic input (control) datasets were available were considered for analysis. To ensure that only the highest quality ChIP-Seq datasets are used for downstream analysis, only those that satisfied the following criteria were retained for further analysis: (i) number of mapped reads is at least 10 million, (ii) reported signal-to-noise ratio (SNR) (Roadmap Epigenomics et al., 2015)—the degree to which reads are concentrated in peaks versus the background—for the H3K4me3 (or H3K27me3) dataset from a given cell/tissue type is (a) greater than that for corresponding genomic input and (b) at least (mean – 1 STD) of SNRs of all H3K4me3 (or H3K27me3, respectively) datasets across all cell/tissue types. (iii) reported SNR for the genomic input from a given cell/tissue type is at most (mean + 1 STD) of SNRS of all input datasets across all cell/tissue types. List of genes DNA-hypermethylated in human osteosarcoma (Easwaran et al., 2012), primary colorectal tumor (Widschwendter et al., 2007), and during aging (Rakyan et al., 2010) were obtained from previously published studies. H3K4me3 and H3K27me3 ChIP-Seq data for osteoblasts and osteosarcoma (Easwaran et al., 2012) were obtained from NCBI GEO portal and aligned to the human genome (hg19) using Bowtie, as described above.

### Data analysis

#### Histone modification enrichment at gene promoters

For each gene promoter, read densities (RPM, reads per million mapped reads) of individual histone modifications (H3K4me3/H3K27me3) and corresponding genomic input were calculated. Promoters were defined as the region spanning TSS ± 500 bp for H3K4me3 and TSS ± 2Kb for H3K27me3. A promoter is deemed to be enriched for a particular histone modification only if its ChIP signal (RPM) is at least (i) 3-fold greater than its input signal (RPM), and (ii) greater than a threshold (1% FDR), estimated as the lowest RPM value at which the number of qualifying promoters (RPM greater than or equal to the threshold) based on the input signal (RPM) is less than 1% of the number of qualifying promoters based on the ChIP signal (RPM). Promoters enriched for both H3K4me3 and H3K27me3 were defined to be bivalent. Promoters enriched for H3K4me3 but not H3K27me3 were defined as H3K4me3-only, and those that are enriched for H3K27me3 but not H3K4me3 were defined as H3K27me3-only. Promoters with neither H3K4me3 nor H3K27me3 enrichment were defined as ‘unmarked’.

#### RNA-Seq data analysis

Reads were aligned to the mouse genome (mm9) using STAR aligner (Dobin et al., 2013), allowing up to three mismatches, retaining only reads that align to unique genomic locations. Alignment files were used to quantitate genes and isoforms annotated in mm9 genome (source: NCBI RefSeq) using the cuffdiff tool (Trapnell et al., 2010), and the resultant “isoforms.fpkm_tracking” file from the cuffdiff run was used to infer differentially expressed genes (*q*-value < 0.05) between two cell-types/time-points of interest.

#### Machine learning approach for predicting bivalent chromatin

Multinomial log-linear models *via* neural networks (multinom function in R) were generated using (a) promoter (±500 bp of TSS) dinucleotide frequencies (5’ to 3’) and (b) promoter (±2Kb of TSS) H3K27me3 tag density (RPM) and enrichment (ChIP/input). A 5-fold cross validation was employed to train and test a model for its ability to classify promoters into one of the four chromatin states: H3K4me3-only, H3K27me3-only, bivalent, and unmarked. At least 1000 models were generated by randomly sampling the dataset for training and testing. Accuracy of a four-class classification was estimated from the confusion matrix for each of the models. Precision and recall for prediction of the bivalent class were estimated from the confusion matrix of the four-class model, where promoters of the bivalent class were considered as positives and those from the other three classes were considered as negatives.

#### Gene promoters hypermethylated in mouse EpiLCs

A gene promoter is defined as hypermethylated in EpiLCs compared to naïve ESCs only if (i) its methylation level in EpiLCs is at least 2-fold greater than that in naïve ESCs, and (ii) its methylation level in EpiLCs is at least 50%.

#### Motif analysis

Occurrences of TF binding motifs within gene promoters (+/- 500 bp of TSS) were inferred using the FIMO tool (Grant et al., 2011). A collection of 746 known motifs from the non-redundant JASPAR CORE 2020 database (Fornes et al., 2020) of vertebrate TF motifs was used as queries. Motif occurrences with p-value < 10^-4^ were deemed significant and were considered for further analysis.

## ACKNOWLEDGMENTS

We thank members of the Jothi Lab for insightful discussions and comments on the manuscript. We thank A.J. Oldfield, J. Rodriguez, and P.A. Wade for critical comments on the manuscript. This work was supported by the Intramural Research Program of the NIH, National Institute of Environmental Health Sciences (1ZIAES102625 to R.J.).

## AUTHOR CONTRIBUTIONS

D.K. and R.J. conceived and designed the study. D.K. performed the research. D.K. and R.J. analyzed the data and wrote the manuscript.

## AUTHOR INFORMATION

The authors declare no competing financial interests. Correspondence and requests for materials should be addressed to R.J. or D.K.

## SUPPLEMENTARY TABLE LEGENDS

**Table S1.** Chromatin states and dinucleotide frequencies of human gene promoters

**Table S2.** Chromatin states and dinucleotide frequencies of mouse gene promoters

**Table S3.** Genes upregulated during ESC to EpiLC differentiation and their temporal gene expression.

**Table S4.** Bivalent genes that harbor poised RNAPII.

**Table S5.** Results from Gene Ontology (GO) analysis.

**Table S6.** Correlation between number of bivalent genes and gene expression.

**Table S7.** Genes hypermethylated in mouse EpiLCs, human osteosarcoma, and human colorectal primary tumor, and human aging.

**Table S8.** Chromatin states of human gene promoters in various cell types.

**Table S9.** Results from sequence motif enrichment analysis.

---

**Figure S1.**
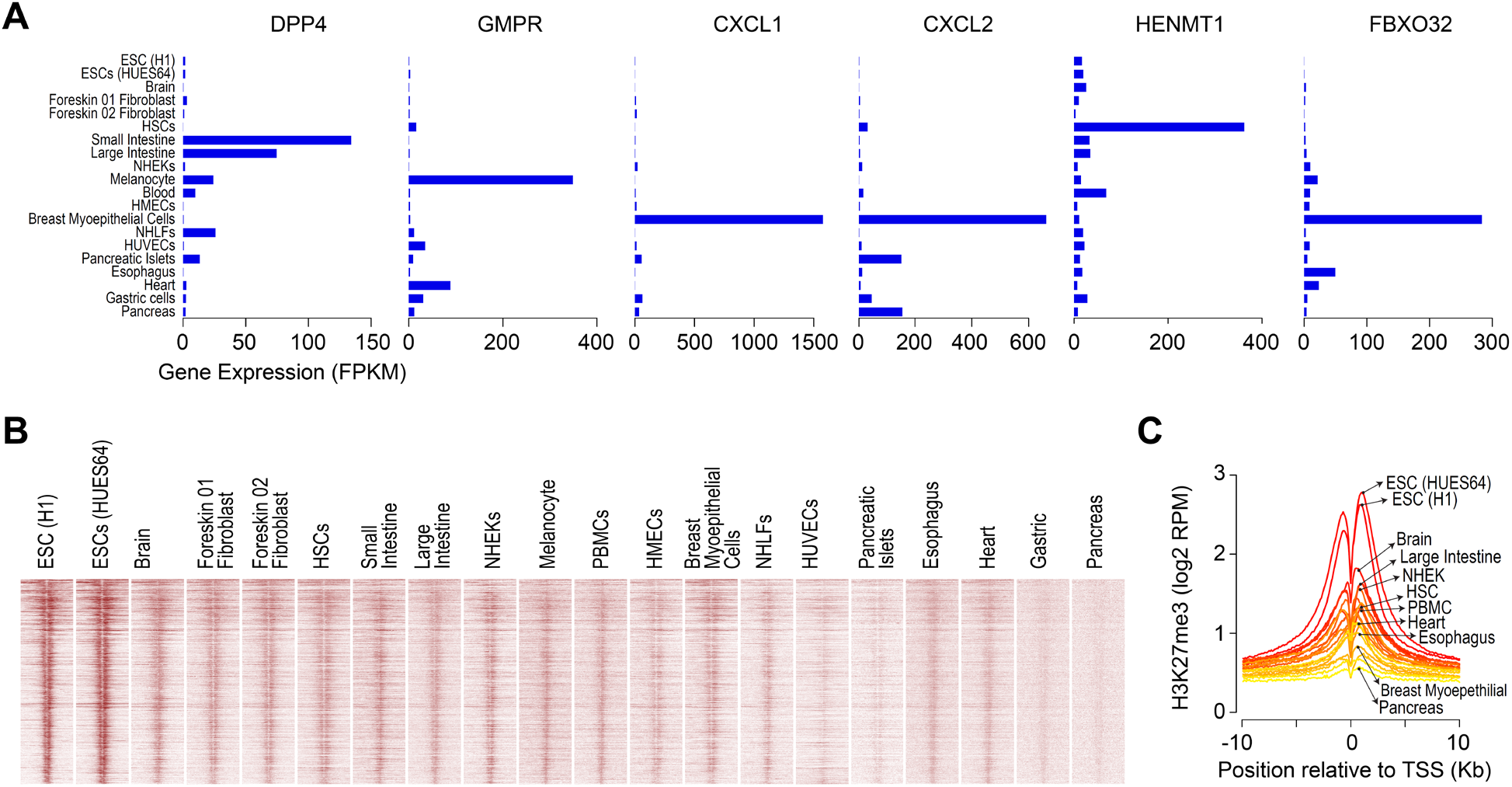
H3K4me3, observed at bivalent promoters in ESCs, persists in nearly all cell types irrespective of gene expression. **(A)** Relative expression of genes, shown in Figure 1A, in various cell types. FPKM, fragments (RNA-Seq) per kilobase per million mapped reads. **(B)** Heatmap representation of H3K27me3 ChIP-Seq read density, in various cell types, near transcription start sites (TSSs) of genes bivalently marked in human ESCs. Ordering of genes (top to bottom) is same as in Figure 1B. Read density is represented as RPM (reads per million mapped reads). **(C)** Average H3K27me3 ChIP-Seq read density, in various cell types, near TSSs of genes shown in B. Shades of color represent individual cell types. Select cell types, out of the twenty plotted, are highlighted.

**Figure S2.**
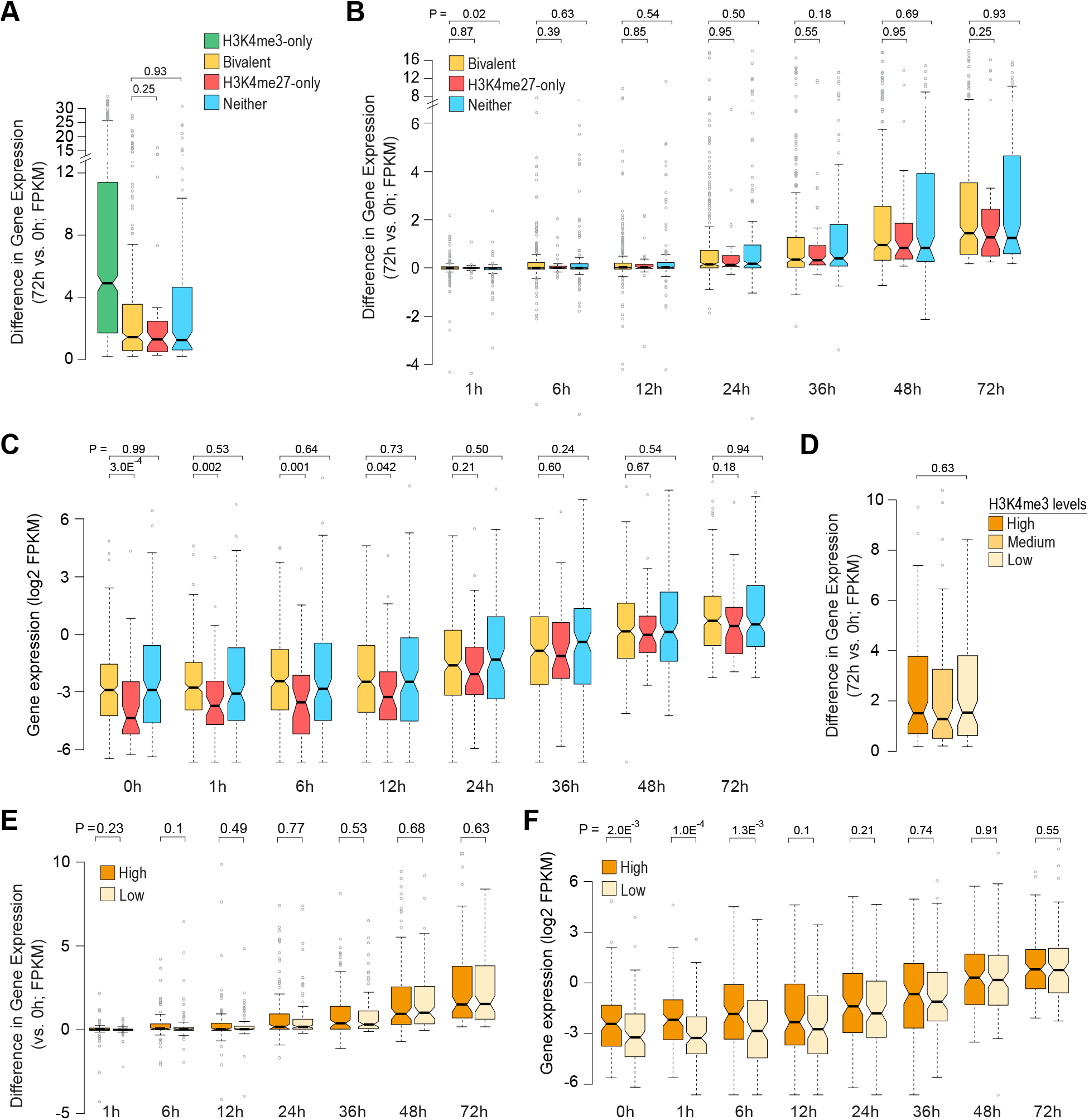
Bivalent chromatin does not poise genes for rapid activation. **(A)** Boxplot showing distribution of absolute differences (delta) in gene expression (72h vs 0h) for genes upregulated in EpiLCs. Genes grouped based on their chromatin states in naïve ESCs (0h). **(B)** Boxplot showing distribution of absolute differences (delta) in gene expression over time (compared to 0h) for genes upregulated in EpiLCs. Genes are grouped based on their chromatin states in naïve ESCs (0h). **(C)** Boxplot showing distribution of absolute gene expression over time for genes upregulated in EpiLCs (72h vs 0h). As in B, genes are grouped based on their chromatin states in naïve ESCs (0h). **(D)** Boxplot showing distribution of absolute differences (delta) in gene expression (72h vs 0h) for bivalent genes upregulated in EpiLCs. Genes are grouped based on H3K4me3 enrichment at promoters in naïve ESCs (0h), as defined in Figure 1H. **(E)** Boxplot showing distribution of absolute differences (delta) in gene expression over time (compared to 0h) for bivalent genes upregulated in EpiLCs. Genes grouped based on high/low H3K4me3 signal, as defined in Figure 1H. **(F)** Boxplot showing distribution of absolute gene expression over time for bivalent genes upregulated in EpiLCs (72h vs 0h). As in E, genes are grouped based on their H3K4me3 levels in naïve ESCs (0h). All the *P* values were calculated using two-sided Wilcoxon rank-sum test.

**Figure S3.**
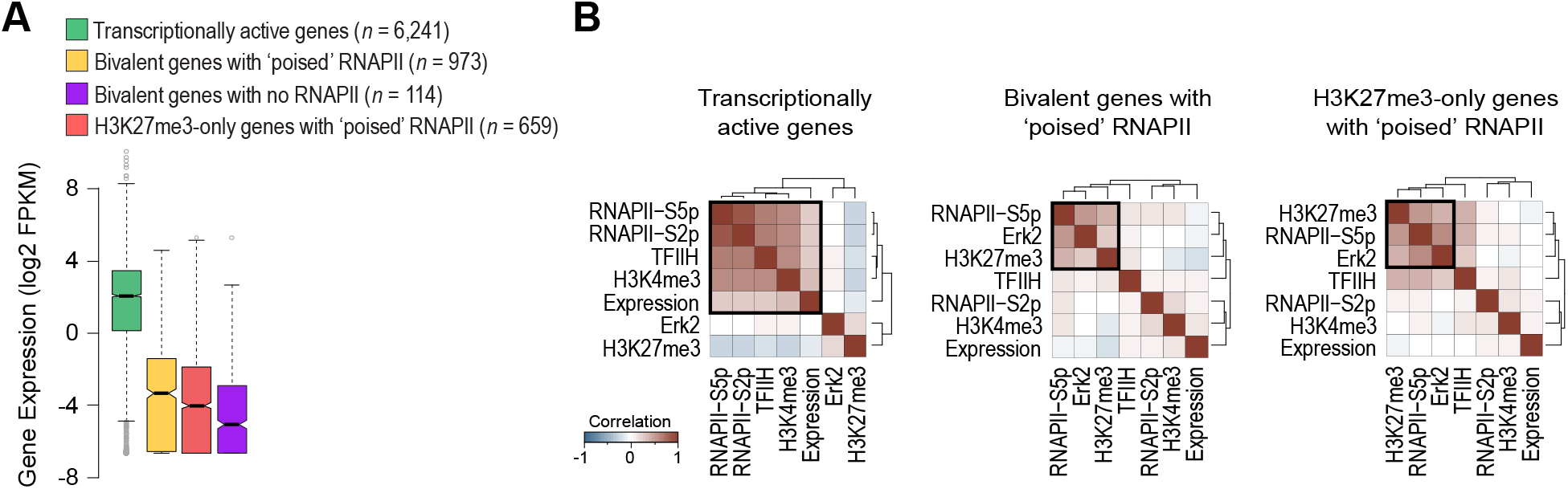
‘Poised’ RNA polymerase II at bivalent genes is incompatible with transcription. **(A)** Boxplot showing the distribution of gene expression in mouse ESCs grown in serum-containing medium (Marks et al., 2012) for the four gene classes shown in Figure 3. FPKM, fragments per kilobase per million mapped reads. **(B)** Heatmaps showing unsupervised hierarchical clustering of pairwise Pearson correlations between signals near TSSs from indicated ChIP-Seq datasets and gene expression (Brookes et al., 2012; Marks et al., 2012; Tee et al., 2014).

**Figure S4.**
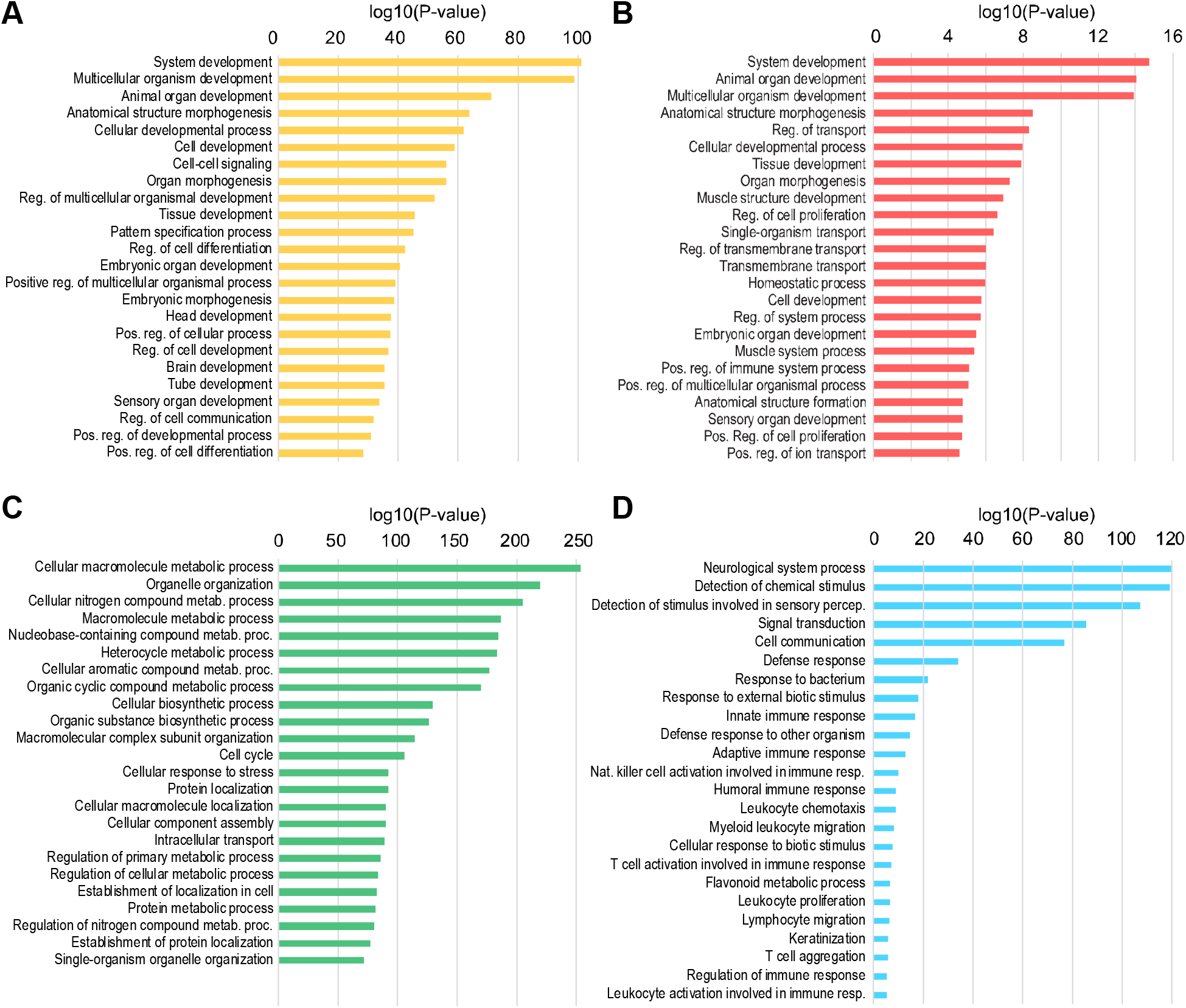
Gene ontology (GO) enrichment analysis. **(A-D)** Top GO categories (biological processes) from enrichment analysis of bivalent (**A**), H3K27me3-only (**B**), H3K4me3-only (**C**), and unmarked (**D**) genes, as defined in naïve mouse ESCs. See **Table S4** for a complete list with additional details.

**Figure S5.**
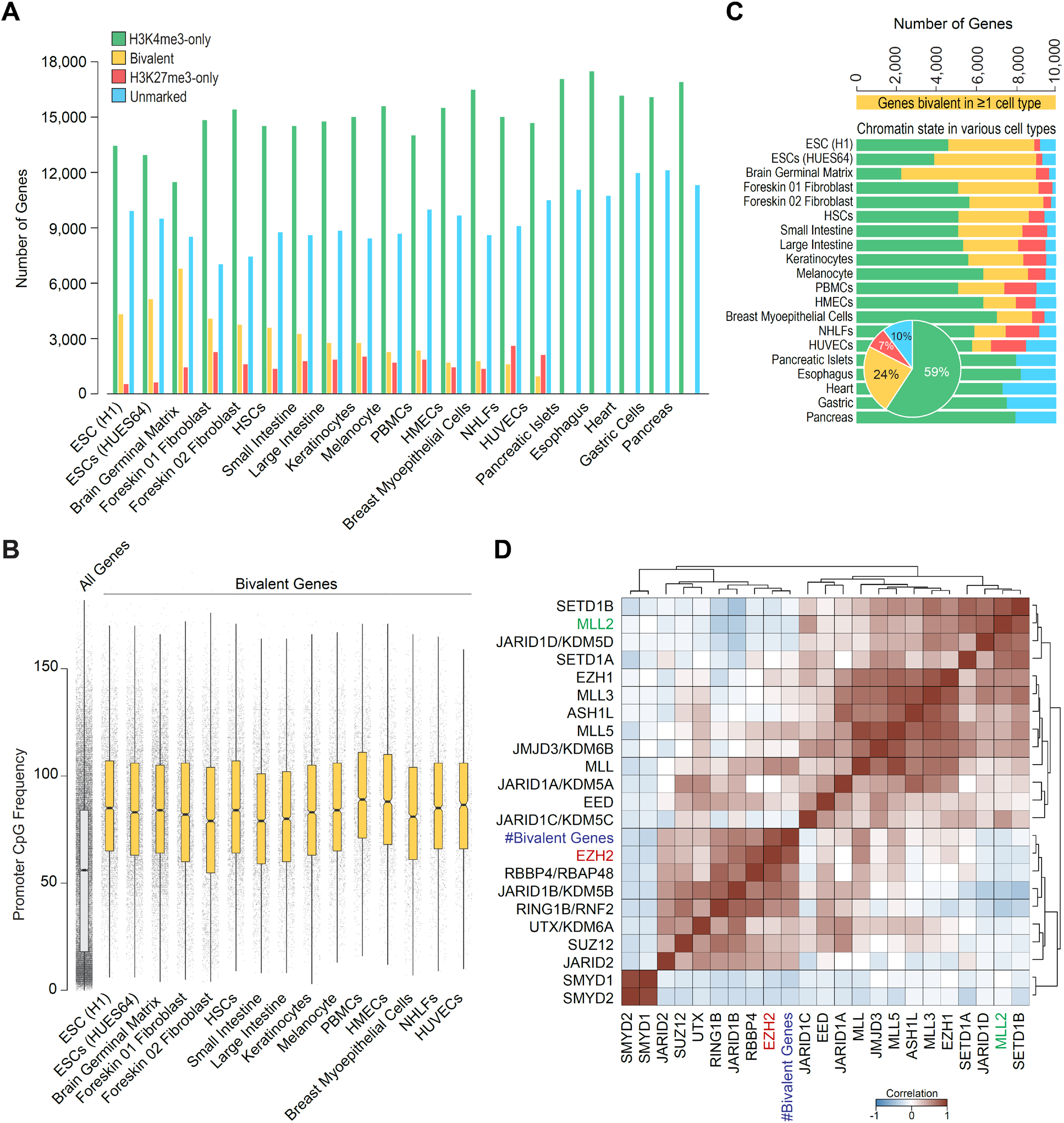
Characterization of bivalent promoters across various cell types. **(A)** Number of genes within each of the four classes, defined based on H3K4me3 (+/-500bp of TSS) and/or H3K27me3 (+/-2Kb of TSS) enrichment at gene promoters in various human cell types. Bivalent, positive for H3K4me3 and H3K27me3; H3K4me3-only, positive for H3K4me3 and negative for H3K27me3; H3K27me3-only, positive for H3K27me3 and negative for H3K4me3; Unmarked, negative for both H3K4me3 and H3K27me3.**(B)** Boxplot showing the distribution of CpG dinucleotide frequency at promoters (+/- 500 bp of TSS) of all genes (left-most) and genes bivalently marked in various human cell types. **(C)** Genes bivalently marked in one or more cell types (*n* = 10,042) and their chromatin state in various cell types (bottom). Inset: pie-chart summarizing the proportional breakdown of chromatin states across all cell types. Color scheme same as in A. **(D)** Heatmap showing unsupervised hierarchical clustering of Pearson correlations between number of bivalent genes and expression levels of H3K4me3 or H3K27me3 methylases and demethylases across cell types.

**Figure S6.**
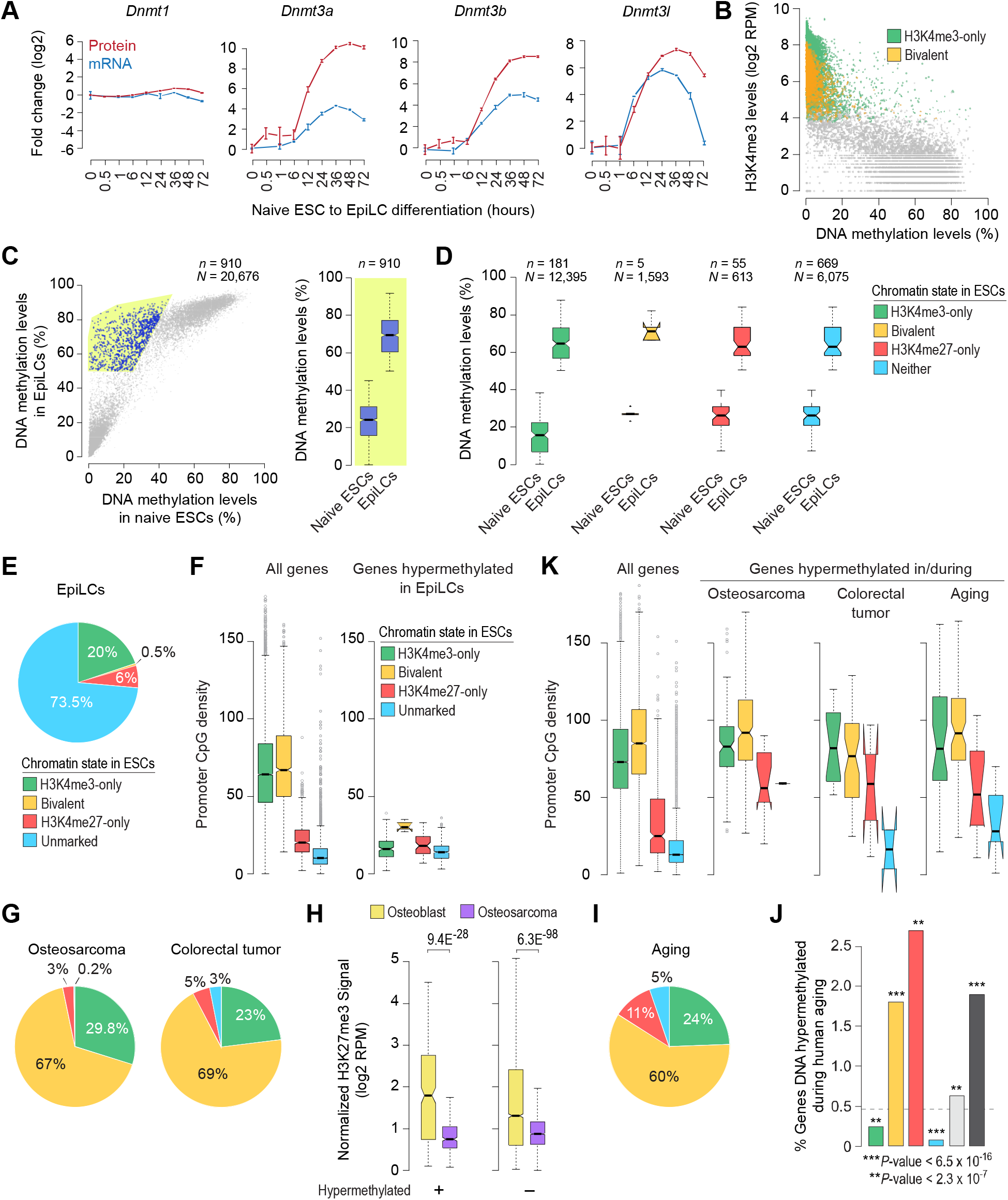
Bivalent chromatin protects promoters from *de novo* DNA methylation. **(A)** Relative protein and mRNA expression levels (compared to 0 h) of DNA methyltransferases (Dnmt1, Dnmt3a, Dnmt3b, Dnmt3l) during naïve mouse ESC to EpiLC differentiation (Yang et al., 2019). Error bars represent SEM. **(B)** Scatter plot showing DNA methylation levels (Shirane et al., 2016) (x-axis) and H3K4me3 levels (Yang et al., 2019) (y-axis) at gene promoters (*N* = 20,676) in naïve mouse ESCs. Individual data points represent gene promoters. H3K4me3-only and bivalent promoters are highlighted in green and orange, respectively. **(C)** *Left*: Scatter plot showing DNA methylation levels at gene promoters (*N* = 20,676) in naïve mouse ESCs (x-axis) and EpiLCs (y-axis) (Shirane et al., 2016). Individual data points represent gene promoters. Promoters that are hypermethylated in EpiLCs compared to ESCs (*n* = 910) are highlighted in blue. *Right*: Boxplot showing the distribution of DNA methylation levels at promoters hypermethylated in EpiLCs compared to ESCs. Methylation data was obtained from **(D)** Promoters hypermethylated in EpiLCs, shown in B, are divided into four groups based on their chromatin state in naïve mouse ESCs. *n*, number of hypermethylated genes within each category; *N*, total number of genes within each category. **(E)** Pie-chart showing proportion of genes hypermethylated in EpiLCs based on their promoter chromatin status in naïve mouse ESCs (Shirane et al., 2016). **(F)** Boxplots showing the distribution of CpG dinucleotide frequency at promoters (+/- 500 bp of TSS) of all genes (left) and those that are hypermethylated in EpiLCs (right). Genes are divided into four groups based on their chromatin state in naïve mouse ESCs. **(G)** Same as in E, but for genes hypermethylated in adult human cancers (Easwaran et al., 2012; Widschwendter et al., 2007). Genes are grouped based on their promoter chromatin status in human ESCs. **(H)** Genes bivalently marked in human ESCs were divided into those that are aberrantly DNA hypermethylated in human osteosarcoma (*left*) and those that are not (*right*). Boxplots show the distribution of H3K27me3 levels at these gene promoters in human osteoblasts (yellow) and osteosarcoma (purple) (Easwaran et al., 2012). **(I)** Same as in G, but for genes DNA hypermethylated during human aging (Rakyan et al., 2010). **(J)** Percentage of genes, within each of the four classes of genes defined in human ESCs, whose promoters are DNA hypermethylated during human aging. Light and dark gray bars respectively denote genes enriched for H3K4me3 (H3K4me3-only and bivalent) and H3K27me3 (bivalent and H3K27me3-only). Dotted gray line denotes expected frequency. **(K)** Same as in F, but for genes hypermethylated in osteosarcoma, colorectal tumor, and during aging. All the *P* values were calculated using two-sided Wilcoxon rank-sum test.

**Figure S7.**
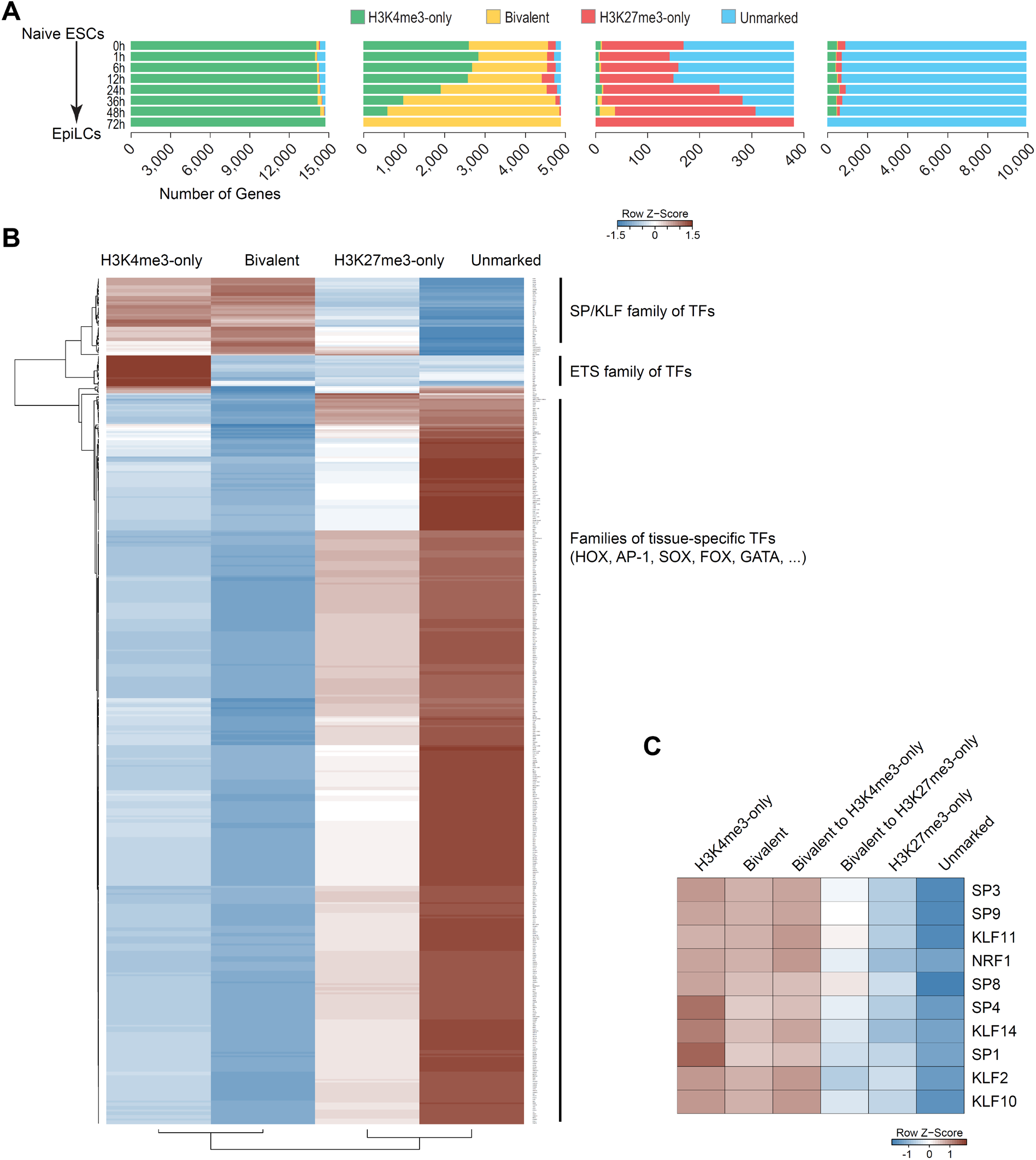
Chromatin fate and sequence characteristics of bivalent promoters. **(A)** Genes are grouped into four classes based on their chromatin state in mouse EpiLCs (72h), defined based on H3K4me3 (+/- 500 bp of TSS) and/or H3K27me3 (+/- 2 Kb of TSS) enrichment at gene promoters, and their chromatin states during mouse ESC to EpiLC transition are shown (top to bottom). **(B)** Heatmap showing relative enrichment for binding motifs for various transcription factors (TFs) within promoters (+/- 500 bp of TSS) of the four genes classes defined based on the chromatin state of promoters in human ESCs. See also **Table S9**. **(C)** Same as in B, for select TFs. Also shown (middle two columns) are relative enrichment within bivalent promoters that mostly resolve into H3K4me3-only or H3K27me3-only state in other cell types.

## Notes

### Competing Interest Statement

The authors have declared no competing interest.

